# Sexual dimorphic regulation of recombination by the synaptonemal complex

**DOI:** 10.1101/2022.10.13.512115

**Authors:** Cori K. Cahoon, Colette M. Richter, Amelia E. Dayton, Diana E. Libuda

## Abstract

In sexually reproducing organisms, germ cells faithfully transmit the genome to the next generation by forming haploid gametes, such as eggs and sperm. Although most meiotic proteins are conserved between eggs and sperm, many aspects of meiosis are sexually dimorphic. The mechanisms regulating recombination display sex-specific differences in multiple organisms such that the same proteins in each sex are being utilized in different ways to produce sexually dimorphic outcomes. The synaptonemal complex (SC), a large ladder-like structure that forms between homologous chromosomes, is essential for regulating meiotic chromosome organization and promoting recombination. To assess whether sex-specific differences in the SC underpin sexually dimorphic aspects of meiosis, we examined two *Caenorhabditis elegans* SC central region proteins, SYP-2 and SYP-3, in oogenesis and spermatogenesis and uncovered sex-specific roles for the SYPs in regulating meiotic recombination. We find that SC composition is regulated by sex-specific mechanisms throughout meiotic prophase I. During pachytene, both oocytes and spermatocytes differentially regulate the stability of SYP-2 and SYP-3 within an assembled SC, with increased SYP-2 dynamics in spermatocytes and increased SYP-3 dynamics in oocytes. Further, we uncover that the relative amount of SYP-2 and SYP-3 within the SC is independently regulated in both a sex-specific and a recombination-dependent manner. Specifically, we find that SYP-2 regulates the early steps of recombination in both sexes, while SYP-3 controls the timing and positioning of crossover recombination events across the genomic landscape in only oocytes. Taken together, we demonstrate dosage-dependent regulation of individual SC components with sex-specific functions in recombination. These sexual dimorphic features of the SC provide insights into how spermatogenesis and oogenesis adapted similar chromosome structures to differentially regulate and execute recombination.

## INTRODUCTION

In most sexually reproducing organisms, meiosis ensures the inheritance of genetic information each generation through the formation of haploid gametes, such as eggs and sperm. Many aspects of meiosis are sexually dimorphic from the differences in the size of egg and sperm cells to the molecular mechanisms ensuring accurate segregation of the chromosomes (reviewed in Cahoon and Libuda 2019). Moreover, many meiotic proteins are present in both sexes. How each sex utilizes a highly similar proteome to produce dimorphic phenotypes remains unclear.

Multiple studies in mice and plants indicate that meiotic chromosome structures, such as the SC, are sexually dimorphic (reviewed in Morgan *et al*. 2017; Cahoon and Libuda 2019). The SC assembles in early prophase I (transition zone/late zygotene) and organizes the genome to both facilitate and enable the essential meiotic processes of homolog pairing and recombination (reviewed in Cahoon and Hawley 2016). The SC forms a scaffold between the homologs that allows for the accurate repair of DNA double strand breaks (DSBs) induced by the topoisomerase-like protein Spo11 (Keeney *et al*. 1997; Zickler and Kleckner 1999; Zickler and Kleckner 2015). A subset of these DSBs must be repaired as crossover recombination events to allow for the accurate segregation of the homologs at anaphase I.

Studies in multiple organisms indicate that the mechanisms that regulate recombination are sexually dimorphic (Cahoon and Libuda 2019). In mice, humans, and some plant species, one sex usually has more crossovers than the other, and the sex with more crossovers typically has longer chromosome axes (Morelli and Cohen 2005; Drouaud *et al*. 2007; Giraut *et al*. 2011; Gruhn *et al*. 2013; Bherer *et al*. 2017; Halldorsson *et al*. 2019). In mice, the placement of crossovers along the chromosomes is different between the sexes. Specifically, recombination hotspots have a sex-bias for crossover formation, which is thought to be driven by sex-specific differences in origin of replication firing and DNA methylation near PRDM9-binding site (Brick *et al*. 2018; Halliwell and Hoffmann 2021; Pratto *et al*. 2021). In *Arabidopsis thaliana*, the positioning and regulation of crossovers are also different between the sexes, with some sex-specific mechanisms, such as epigenetic regulation of recombination, being conserved between mice and plants (Giraut *et al*. 2011; Clement and Arndt 2013; Arbeithuber *et al*. 2015; de Boer *et al*. 2015; Choi *et al*. 2018; Underwood *et al*. 2018). However, some organisms, such as *C. elegans*, lack identified differences in the epigenetic regulation of recombination on the autosomes, but still exhibit sex-specific recombination mechanisms. In *C. elegans*, the timing of recombination initiation and organization of the chromatin is different between the sexes, which together influences the sex-specific mechanisms regulating crossover establishment and chromosome segregation (Jaramillo-Lambert *et al*. 2007; Jaramillo-Lambert *et al*. 2016; Li *et al*. 2020; Rourke and Jaramillo-Lambert 2022). Specifically, in *C. elegans*, the recombination promoting protein BRC-1 and its binding partner the E3 ligase BRD-1 act on different steps of recombination in each sex likely through sex-specific differences in protein interactions and ubiquination of target proteins (Li *et al*. 2020). In *Drosophila melanogaster*, spermatocytes do not require recombination nor the SC to properly segregate homologs, but recombination and the SC are required for proper homolog segregation in oocytes (Page and Hawley 2001; Jeffress *et al*. 2007; Page *et al*. 2008; McKee *et al*. 2012; Collins *et al*. 2014). Although many sexual dimorphisms have been discovered in multiple organisms, how each sex uses the same or similar proteomes to differentially influence recombination outcomes remains unclear.

The SC is both required for the formation of crossovers and influences the frequency of crossovers occurring per homolog pair in multiple organisms (Sym *et al*. 1993; Yuan *et al*. 2000; Page and Hawley 2001; MacQueen *et al*. 2002; Yuan *et al*. 2002; Colaiacovo *et al*. 2003; Hillers and Villeneuve 2003; Anderson *et al*. 2005; Costa *et al*. 2005; de Vries *et al*. 2005; Higgins *et al*. 2005; Bolcun-Filas *et al*. 2007; Jeffress *et al*. 2007; Smolikov *et al*. 2007a; Hamer *et al*. 2008; Page *et al*. 2008; Smolikov *et al*. 2009; Hayashi *et al*. 2010; Schramm *et al*. 2011; Yuan *et al*. 2012; Libuda *et al*. 2013; Collins *et al*. 2014; Woglar and Villeneuve 2018; Hurlock *et al*. 2020; Gordon *et al*. 2021; Liu *et al*. 2021). Specific proteins within the SC can influence crossover distribution, DSB repair mechanisms, and specific steps in the crossover licensing process (Jeffress *et al*. 2007; Voelkel-Meiman *et al*. 2015; Voelkel-Meiman *et al*. 2019; Gordon *et al*. 2021; Lascarez-Lagunas *et al*. 2022; Voelkel-Meiman *et al*. 2022). In *C. elegans*, pro-crossover proteins are recruited to the SC by the central region (SYP) proteins of the SC (Libuda *et al*. 2013; Cahoon *et al*. 2019). The exact mechanism(s) of how the SC regulates crossing over remains an active area of study in multiple organisms. Notably, the sexual dimorphisms present in the mechanisms regulating DSB repair suggest that SC-dependent regulation of crossing over may also be sexually dimorphic (Brick *et al*. 2018; Li *et al*. 2020). Further, the SC in multiple organisms has sex-specific requirements with some SC mutants displaying sex-specific fertility defects that are limited to one sex (reviewed in Morelli and Cohen 2005; Cahoon and Libuda 2019). Taken together, the sexually dimorphic regulation of recombination and sexually dimorphic properties of SC mutants suggests that the composition and/or function of the SC is different between oogenesis and spermatogenesis.

Similar to other organisms, many aspects *C. elegans* meiosis is sexually dimorphic from the timing of egg and sperm development to the regulation of checkpoints and recombination (Gumienny *et al*. 1999; Jaramillo-Lambert *et al*. 2007; Jaramillo-Lambert and Engebrecht 2010; Jaramillo-Lambert *et al*. 2010; Lamelza and Bhalla 2012; Checchi *et al*. 2014; Jaramillo-Lambert *et al*. 2016; Saito and Colaiacovo 2017; Li *et al*. 2020; Cahoon and Libuda 2021; Gartner and Engebrecht 2022; Rourke and Jaramillo-Lambert 2022). These sex-specific differences suggest that other critical recombination regulatory processes, such as the SC, may also have sexually dimorphic features in *C. elegans*. Both sexes in *C. elegans* are assumed to assemble the same SYP proteins into the SC, but this aspect has not been extensively investigated.

The SC structure in *C. elegans* can be divided into two parts: lateral elements and the central region (Figure 1A). The lateral element proteins assemble along the chromosome axis of each homolog and the central region proteins span the gap between the homologous chromosomes. In *C. elegans*, the central region proteins of the SC are known as SYP proteins. To date, six SYP proteins have been identified (SYP-1 through SYP-6) (MacQueen *et al*. 2002; Colaiacovo *et al*. 2003; Smolikov *et al*. 2007a; Smolikov *et al*. 2009; Hurlock *et al*. 2020; Liu*et al*. 2021). The most well characterized central region SC proteins are SYP-1, SYP-2, and SYP-3 (Figure 1A). SYP-1, the transverse filament protein, spans the distance between the homologs and the N-terminus is important for crossover regulation (MacQueen *et al*. 2002; Schild-Prufert *et al*. 2011; KÖhler *et al*. 2020; Gordon *et al*. 2021). SYP-2 is located as a single track down the middle of the SC and phosphorylation of SYP-2 is involved in regulating the timing of SC disassembly (Colaiacovo *et al*. 2003; Schild-Prufert *et al*. 2011; Nadarajan *et al*. 2016). SYP-3 is positioned broadly within the central region of the SC as two parallel tracks surrounding the single SYP-2 track in the middle (Smolikov *et al*. 2007b; Schild-Prufert *et al*. 2011). In oocytes, SYP-3 is highly dynamic during pachytene but becomes progressively stabilized within the SC as crossovers are established (Nadarajan *et al*. 2017; Pattabiraman *et al*. 2017). While the lateral element proteins do not display the same dynamics as SYP-3 (Rog *et al*. 2017), it remains unknown whether other SYP proteins display these same properties as SYP-3. Further, most of our knowledge about the SC in *C. elegans* has been found using oocytes, thus any spermatocyte-specific roles for the SC remain unknown.

**Figure 1:**
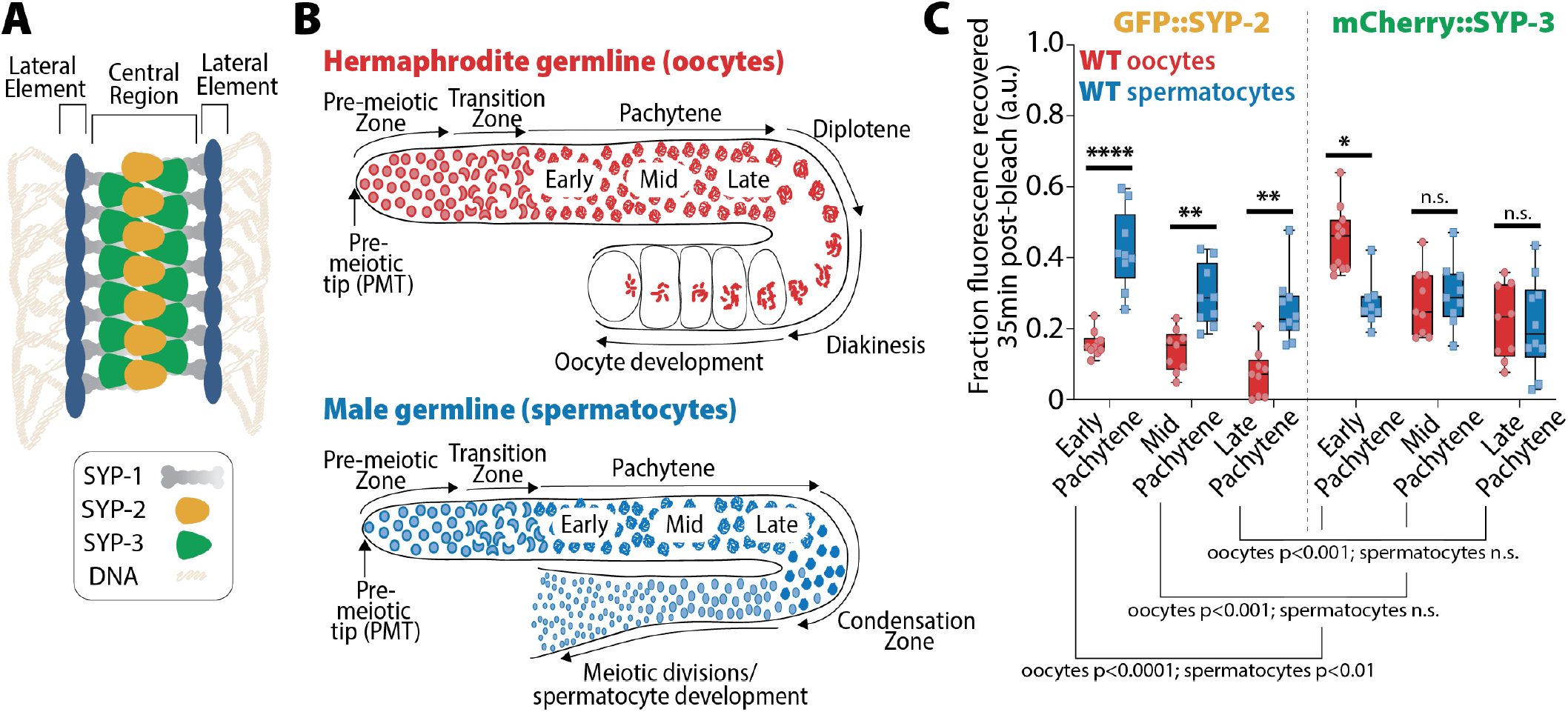
SYP-2 and SYP-3 dynamics are sexually dimorphic. **(A)** Diagram of the synaptonemal complex showing the locations SYP-1 (gray), SYP-2 (yellow), and SYP-3 (green) within the central region of the complex. **(B)** Diagrams of hermaphrodite (top, red) and male (bottom, blue) germlines with developing oocytes and spermatocytes, respectively. The stages of the germlines are labeled starting at the pre-meiotic tip (PMT) and ending at the meiotic divisions. **(C)** Quantification of the fraction of fluorescence recovered 35 minutes after photobleaching a small region of either GFP::SYP-2 (left) or mCherry::SYP-3 (right) in both oocytes (red)and spermatocytes (blue) (a.u. = arbitrary units). All statistics are multiple comparisons adjusted Mann Whitney U test and unless the p value is indicated the asterisk indicates number of significant digits from p=0.05 (n.s. = not significant). Oocyte data with GFP::SYP-2 is from 10 nuclei (early), 9 nuclei (mid), and 9 nuclei (late), and with mCherry::SYP-3 is from 11 nuclei (early), 9 nuclei (mid), and 9 nuclei (late). Spermatocyte data with GFP::SYP-2 is from 9 nuclei (early), 9 nuclei (mid), and 10 nuclei (late), and with mCherry::SYP-3 is from 8 nuclei (early), 9 nuclei (mid), and 10 nuclei (late).

Here we show that the SC is sexually dimorphic in *C. elegans*. Specifically, we demonstrate that the composition of the SC is not uniform during prophase I and instead is regulated in a sex-specific and protein dosage-dependent manner to facilitate specific steps of recombination. We find that SYP-2 dosage is critical for the establishment and/or stabilization of recombination intermediates, while SYP-3 dosage modulates the timing of crossover designation during pachytene. Taken together, our study reveals novel regulation of recombination whereby the SC composition is dynamically altered throughout pachytene to facilitate sexually dimorphic mechanisms of DNA repair.

## RESULTS

### SYP-2 and SYP-3 are sexually dimorphic

To understand the relationship between the sexual dimorphic aspects of recombination and the SC, we used the model system *C. elegans* where oocyte and spermatocyte development can be easily accessed and analyzed at both the same time and developmental stage of the animal. Adult male worms undergo spermatogenesis, while adult hermaphrodite worms undergo oogenesis (Figure 1B). The germline for both sexes is organized as a spatial-temporal gradient along the distal-proximal axis, thereby allowing for easy access to all stages of meiotic prophase I simultaneously (Hillers *et al*. 2017; Gartner and Engebrecht 2022). Nuclei proliferate at the distal end of the germline (pre-meiotic tip) and physically move proximately as they proceed into the stages of meiosis: the transition zone (leptotene/zygotene), pachytene, diplotene, and diakinesis (in spermatocytes diplotene/diakinesis is termed the condensation zone). At the end of prophase I, oocytes nuclei arrest at diakinesis until they are fertilized, but spermatocytes rapidly complete the meiosis I and meiosis II divisions to generate mature sperm.

To assess if SC dynamics differ between spermatocytes and oocytes, we performed fluorescent recovery after photobleaching (FRAP) assays with two SC proteins endogenously tagged with fluorescent proteins: (1) GFP::SYP-2 from (Gao *et al*. 2016), and (2) mCherry::SYP-3 that we generated using CRISPR/Cas9 (see Methods) (Figure 1C, S1). We found that both SYP-2 and SYP-3 have sex-specific differences in protein turnover, with SYP-2 dynamics higher in spermatocytes and SYP-3 dynamics higher in oocytes.

For GFP::SYP-2, spermatocytes displayed significantly higher recovery dynamics throughout pachytene compared to oocytes (Figure 1C, S1; early pachytene oocyte 15.8% vs. spermatocyte 42.2% mean recovery, P<0.0001; mid pachytene oocyte 13.8% vs. spermatocyte 29.3% mean recovery, P=0.0009; oocyte 7.4% vs spermatocyte 25.3% mean recovery late pachytene P=0.0002; Bonferroni-Dunn adjusted, Mann-Whitney). In contrast, mCherry::SYP-3 recovered more quickly in oocytes during early pachytene compared to spermatocytes (early pachytene oocyte 45.1% vs. spermatocyte 27.2% mean recovery, P=0.0015; Bonferroni-Dunn adjusted, Mann-Whitney). SYP-3 recovery rates were similar between the sexes during mid and late pachytene (mid pachytene oocyte 27.5% vs. spermatocyte 29.6% mean recovery, P>0.9999; late pachytene oocyte 21.6% vs. spermatocyte 21.0% mean recovery, P>0.9999; Bonferroni-Dunn adjusted, Mann-Whitney). Both sexes showed the same overall trend of progressive stabilization of SYP-2 and SYP-3 throughout pachytene, which matches previous observations with the transgene GFP::SYP-3 (Nadarajan *et al*. 2017; Pattabiraman *et al*. 2017; Rog *et al*. 2017). Notably, we observed that the progressive stabilization of spermatocyte SYP-3 was less pronounced and much more subtle than that of oocyte SYP-3, suggesting that spermatocytes may not modulate SYP-3 stability via the same mechanisms as oocytes (Figure 1C). Collectively, these data indicate that SYP dynamics are not uniformly regulated throughout prophase and instead exhibit both SYP-specific and sex-specific dynamics.

Upon comparing SYP mobilization between the sexes, we found that SYP-2 and SYP-3 displayed differences in mobility when compared to each other within sex. In oocytes, GFP::SYP-2 turnover was significantly reduced compared to mCherry::SYP-3 turnover (Figure 1C, S1; early pachytene oocyte SYP-2 15.8% vs oocyte SYP-3 45.1% mean recovery, P<0.0001; mid pachytene oocyte SYP-2 13.8% vs oocyte SYP-3 27.5% mean recovery, P=0.0037; late pachytene oocyte SYP-2 7.4% vs oocyte SYP-3 21.6% mean recovery P=0.0083; Bonferroni-Dunn adjusted, Mann-Whitney). Comparatively in spermatocytes, only early pachytene displayed significant differences in GFP::SYP-2 and mCherry::SYP-3 turnover (Figure 1C, S1; early pachytene spermatocyte SYP-2 42.2% vs spermatocyte SYP-3 27.2% mean recovery P=0.0237, Bonferroni-Dunn adjusted, Mann-Whitney). These results demonstrate that SYP-2 and SYP-3 can be independently regulated during pachytene.

To determine whether differential regulation of SYP-2 and SYP-3 might be influenced by chromosome length, we measured SC length during early, mid, and late pachytene in both sexes. Our results found that SC length was similar throughout pachytene in each sex (Figure S2B). The slightly shorter SC lengths in late pachytene spermatocytes was likely due to differences in chromosome compaction between the sexes (Samson *et al*. 2014; Rourke and Jaramillo-Lambert 2022). These results suggest that independent regulation of SYP-2 and SYP-3 within an assembled SC is not due to changes in chromosome length, but rather by other factors.

### SYP-2 and SYP-3 composition within the SC is sexually dimorphic

The differences in SYP-2 and SYP-3 mobility between oogenesis and spermatogenesis suggests that the abundance or concentration of each protein within the SC may also be sexually dimorphic. To compare the relative SYP compositions within the SC between sexes during meiotic prophase I, we calculated the mean fluorescence intensity per cubic micrometer of SC for individual nuclei throughout pachytene in oogenesis and spermatogenesis, and then calculated the average SYP intensity among nuclei in a sliding window across the normalized germline length (see Methods). We found that the accumulation of GFP::SYP-2 in the SC is sexually dimorphic (Figure 2A, 2B, Table S1). Wild type oocytes progressively accumulated SYP-2 up until the early to mid pachytene transition (early pachytene mean intensity 153,292.27 ± SD 13,983.85; early/mid pachytene mean peak intensity 161,878.14 ± SD 13,930.43; mid pachytene mean intensity 148,430.1 ± SD 13,058.02) and then reduce the amount of SYP-2 slightly before maintaining a relatively constant level of SYP-2 through late pachytene (late pachytene mean intensity 143,165.9 ± SD 10,510.79). In contrast to oocytes, wild type spermatocytes progressively loaded SYP-2 into the SC until the onset of late pachytene at which point the amount of SYP-2 began to stabilize (Figure 2A, 2B, Table S1). Notably, early pachytene spermatocytes had less SYP-2 loaded than oocytes (oocytes mean intensity 153,292.27 ± SD 13,983.85 vs. spermatocytes mean intensity 120,002.07 ± SD 20,899.17, P<0.001, Bonferroni adjusted, Mann-Whitney), but late pachytene spermatocytes had near or slightly more amounts of SYP-2 in the SC than oocytes (oocytes mean intensity 143,165.9 ± SD 10,510.79 vs. spermatocytes mean intensity 148,742.2 ± SD 17,003.32, P<0.001, Bonferroni adjusted, Mann-Whitney). Additionally, slightly past the early/mid pachytene transition of SYP-2 intensity, we noted only oocytes display a few persisting bright nuclei in nearly all germlines examined (8 out of 9 germlines with persisting bright nuclei; Figure 2B). Thus, the incorporation of SYP-2 throughout pachytene differs between sexes.

**Figure 2:**
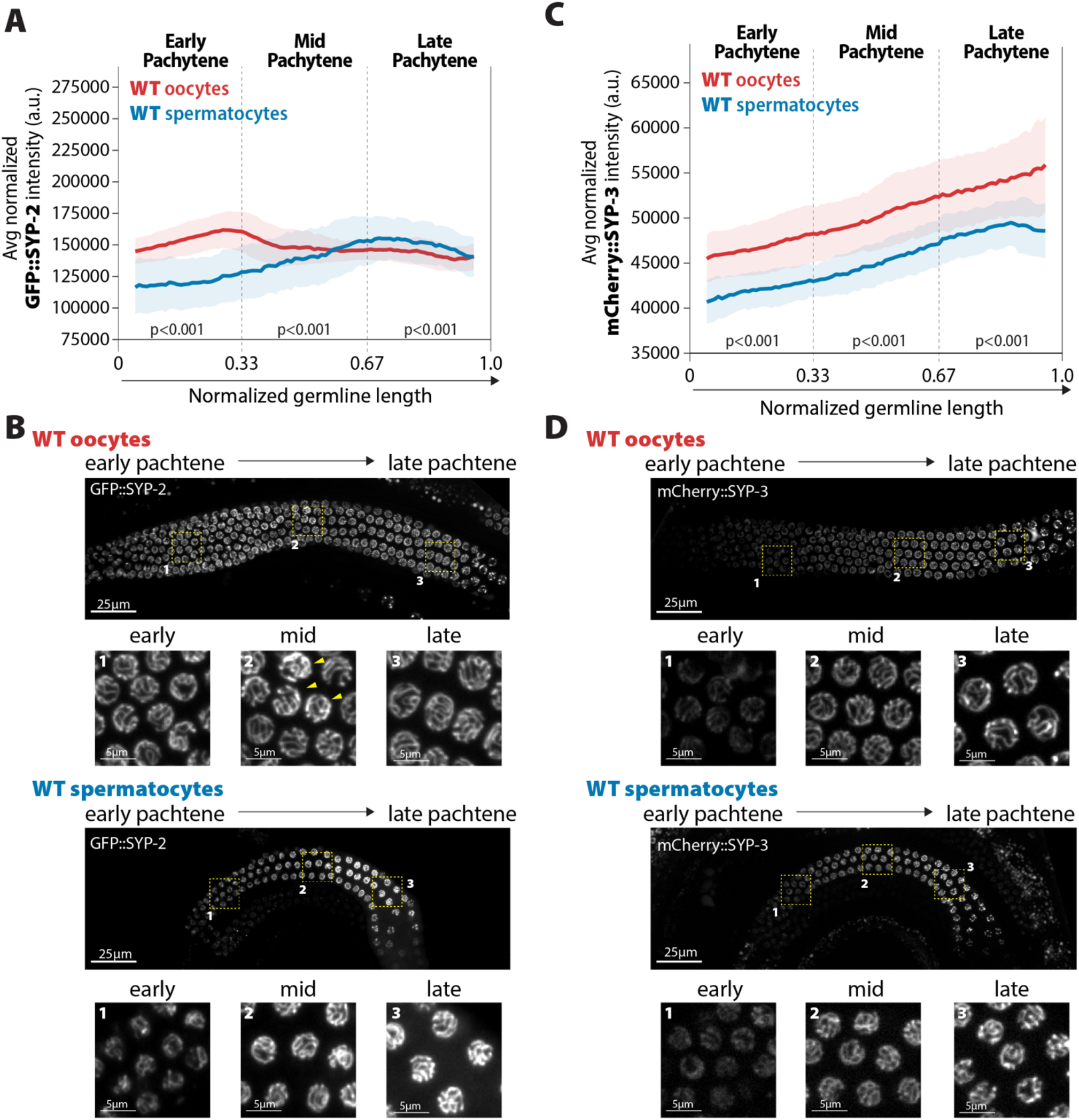
Accumulation of SYP-2 and SYP-3 in the SC are sexually dimorphic. **(A**,**C)**Quantification of the mean intensity of GFP::SYP-2 **(A)** or mCherry::SYP-3 **(C)** per nucleus normalized by the volume of each nucleus (see Methods) throughout pachytene for wild type (WT) oocytes (red, pale red band is the standard deviation) and spermatocytes (blue, pale blue band is the standard deviation). P values on the plot are comparisons between oocytes and spermatocytes for each region of pachytene using Mann Whitney U tests. **(B**,**D)** Represented images of GFP::SYP-2 **(B)** or mCherry::SYP-3 **(D)** in wild type (WT) hermaphrodite (top, oocytes) and male (bottom, spermatocytes) germlines. Germlines are oriented with the start of pachytene on the left and meiotic progression continues to the right. Yellow boxes identify the regions enlarged in each image below to show representatives of early, mid and late pachytene. Arrowheads indicate GFP::SYP-2 bright nuclei in mid pachytene. The intensity adjusts are the same for both GFP::SYP-2 germlines and mCherry::SYP-3 germlines, respectively. n values for number of germlines and nuclei can be found in Table S1.

In contrast to SYP-2, mCherry::SYP-3 progressively accumulated throughout pachytene in both sexes, matching previous observations using a GFP::SYP-3 transgene (Figure 2C, 2D, Table S1) (Pattabiraman *et al*. 2017). Spermatocytes incorporated less SYP-3 throughout pachytene compared to oocytes (early pachytene oocytes mean intensity 46,702.59 ± SD 3047.98 vs. spermatocytes mean intensity 41,820.06 ± SD 2234.53, P<0.001; mid pachytene oocytes mean intensity 50,134.82 ± SD 3715.01 vs. spermatocytes mean intensity 45,021.34 ± SD 2654.22, P<0.001; late pachytene oocytes mean intensity 54,089.69 ± SD 4487.15 vs. spermatocytes mean intensity 48,640.05 ± SD 2727.99, P<0.001; Bonferroni adjusted, Mann-Whitney), which was also similarly observed in the early regions of pachytene with SYP-2. Thus, spermatocytes incorporate less SYP-3 and SYP-2 in the SC than oocytes, specifically within the early and mid regions of pachytene. Moreover, unlike with SYP-2, we did not observe in either sex any bright SYP-3 nuclei that were not surrounded by other nuclei of similar intensity, thereby suggesting that these bright SYP-2 nuclei may have defects or changes that only trigger a response with SYP-2 levels (Figure 2D, Table S1). Taken together, these data demonstrate that SYP-2 and SYP-3 are differentially incorporated in the SC both over the course of meiotic prophase I and between sexes.

### Sex-specific recombination dependent regulation SYP-2 and SYP-3 within the SC

During pachytene, one of the main functions of the SC is to facilitate and regulate recombination. In *C. elegans* oocytes, SPO-11 induced DSBs are formed in the context of fully assembled SC in early pachytene, and these breaks get repaired as the nuclei traverse through pachytene (reviewed in Gartner and Engebrecht 2022). By the transition to late pachytene, crossover recombination events are designated and marked by the pro-crossover protein COSA-1 (Yokoo *et al*. 2012). To determine if the changes we observed in SYP-2 and SYP-3 accumulation during pachytene were caused by recombination, we assessed how the absence of recombination influenced the incorporation of each protein within the SC. To achieve this, we inhibited the formation of crossovers at different stages of recombination using two mutants: (1) *spo-11(me44)* mutants, which cannot form DSBs; and (2) *cosa-1(tm3298)* mutants, which cannot designate DSBs for crossover formation.

The amount of GFP::SYP-2 loaded into the SC was significantly increased in *spo-11* mutant oocytes, but not spermatocytes (Figure 3A, Table S1). Further, the pattern of SYP-2 accumulation was altered in *spo-11* oocytes, in which SYP-2 amounts continued to increase within the SC displaying a 1.27-fold increase in SYP2 amounts during early pachytene, 1.62-fold increase during mid pachytene, and 1.57-fold increase during late pachytene (Figure 3A solid red line compared to dashed black line; early pachytene: mean intensity wild type 153,292.27 ± SD 13,983.85 vs. *spo-11* 195304.87 ± SD 24,775.95; mid pachytene: mean intensity wild type 148,430.1 ± SD 13058.02 vs. *spo-11* 240,817.2 ± SD 31,477.76; late pachytene: mean intensity wild type 143,165.9 ± SD 10,510.79 vs. *spo-11* 224,279.3 ± SD 43,720.01). In contrast, *spo-11* spermatocytes displayed mild changes in amounts of SYP-2 in the SC during pachytene compared to wild type (Figure 3A, solid red line compared to solid blue line). Additionally, the overall trend of SYP-2 incorporation remains unaltered in *spo-11* spermatocytes (Figure 3A solid blue line compared to dashed gray line; early pachytene 0.88-fold decrease in SYP-2 amount; mid pachytene 0.92-fold decrease in SYP-2 amount; late pachytene 0.97-fold decrease in SYP-2 amount). Thus, oocytes and spermatocytes respond in distinctly different ways when DSB formation is eliminated, demonstrating that the incorporation of SYPs is regulated in a recombination-dependent and in a sex-specific manner during prophase I.

**Figure 3:**
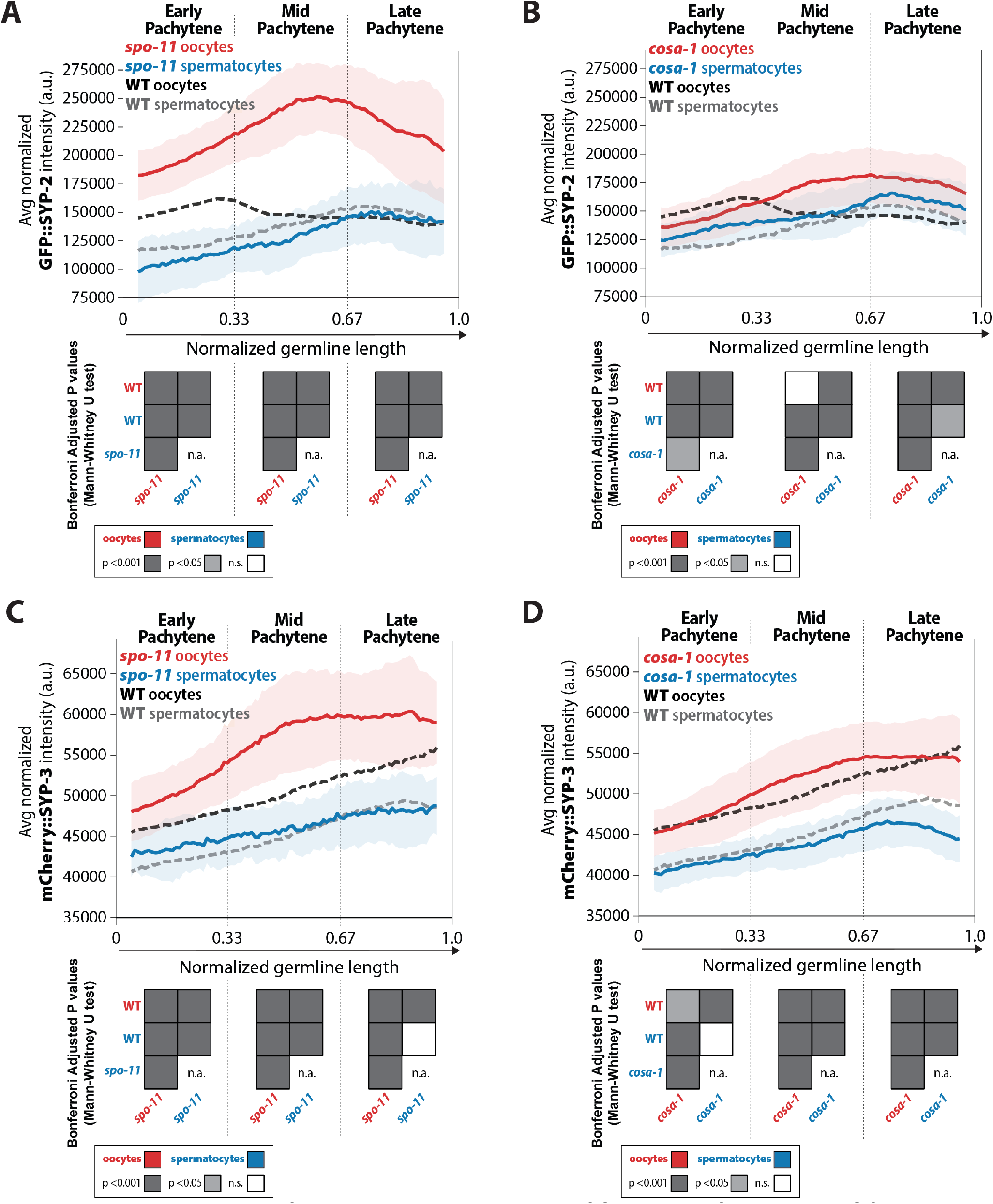
Recombination influences the incorporation of SYP-2 and SYP-3 in the SC differently in each sex. **(A**,**B)** Quantification of the mean intensity of GFP::SYP-2 per nucleus normalized by the volume of each nucleus (see Methods) throughout pachytene for *spo-11* **(A)** and *cosa-1* **(B). (C**,**D)** Quantification of the mean intensity of mCherry::SYP-3 per nucleus normalized by the volume of each nucleus (see Methods) throughout pachytene for *spo-11* **(C)** and *cosa-1* **(D)**. Oocytes are shown in red with the standard deviation shown as a pale red band and spermatocytes are shown in blue with the standard deviation shown as a pale blue band. The mean intensity of GFP::SYP-2 **(A**,**B)** or mCherry::SYP-3 **(C**,**D)** for wild type (WT) oocytes (black) and wild type (WT) spermatocytes (gray) are shown as dashed lines. Heat maps below each pachytene region show the Bonferroni adjusted P values from Mann-Whitney U tests with dark gray indicating p<0.001, light gray indicating p<0.05, and white indicating not significant (n.s.). The self-comparison between spermatocyte *spo-11* or *cosa-1* mutants was not determined (n.a.). n values for number of germlines and nuclei can be found in Table S1.

This sex-specific regulation of the SYPs in response to DSB formation was also observed in the crossover deficient *cosa-1* mutants, albeit to a different degree in comparison to *spo-11* mutant oocytes (Figure 3B, 3C, Table S1). Specifically, *cosa-1* oocytes did not increase the levels of SYP-2 and SYP-3 to the same amounts as observed in *spo-11* oocytes (Figure 3, Table S1). In early pachytene, *cosa-1* oocytes showed a 0.8-fold decrease in SYP-2 amounts that changed in mid and late pachytene with SYP-2 amounts increasing by 1.16-fold and 1.22-fold over wild type amounts, respectively (Figure 3C, solid red line vs. dashed black line). Whereas *spo-11* oocytes increased the amount of assembled SYP-2 to a greater degree than *cosa-1* oocytes having SYP-2 amounts increase by 1.27-fold in early pachytene, 1.62-fold in mid pachytene, and 1.57-fold in late pachytene (Figure 3B, solid red line vs. dashed black line). Thus, SYP-2 and SYP-3 levels in oocytes can be differentially regulated depending on the specific recombination stage that is impeded. Similar to *spo-11* mutant spermatocytes, *cosa-1* mutant spermatocytes did not largely alter SYP-2 and SYP-3 levels (Figure 3, Table S1). Even when recombination is hindered, the pattern of SYP-2 and SYP-3 incorporation throughout pachytene is not the same for each sex or between the SYPs. Overall, the SYPs within the SC of oocytes largely respond to alterations in recombination whereas the SC of spermatocytes do not significantly respond to recombination.

### Reduced SYP-2 causes altered SC assembly in oocytes

The SYP-specific changes in SC composition and in response to recombination defects raised the possibility that SYP protein dosage may regulate specific steps of recombination. To alter the dosage of each SYP, we used heterozygous null mutants for either *syp-2(ok307)* or *syp-3(ok785)* referred to as *syp-2*/+ or *syp-3*/+. In oocytes, reducing the dosage of SYP-1, SYP-2 or SYP-3 by 60-70% was sufficient to permit SC assembly and crossover designation (Libuda *et al*. 2013). Although it remained unclear whether altering the dosage of the SYPs also influenced chromosome pairing or the timing of SC assembly, which can also influence downstream meiotic processes like recombination (Couteau *et al*. 2004; Nabeshima *et al*. 2004; Couteau and Zetka 2005; MacQueen *et al*. 2005; Martinez-Perez and Villeneuve 2005; Goodyer *et al*. 2008; Zhang *et al*. 2012; Mlynarczyk-Evans and Villeneuve 2017). In *syp-2*/+ and *syp-3*/+ mutants, the transition zone (determined by DAPI-stained DNA morphology) was not significantly extended in either sex, indicating that chromosome pairing is not impaired by SYP dosage (Figure S3). Since SYP-1, SYP-2 and SYP-3 are dependent on each other for assembly, we assessed SC assembly using SYP-1 staining in *syp-2*/+ and *syp-3*/+ oocytes and spermatocytes (MacQueen *et al*. 2002; Colaiacovo *et al*. 2003; Smolikov *et al*. 2007b). SC assembly and/or the maintenance of full length SC was altered by reducing the dosage of SYP-2 in only oocytes (Figure 4). In early pachytene, *syp-2*/+ oocytes displayed more discontinuities in the SC along the chromosomes than both wild type and *syp-3*/+ suggesting that there is a SC assembly or maintenance defect in *syp-2*/+ (Figure 4A, yellow arrow heads). Notably, these SC defects in early pachytene caused a significant increase in the length of the SC assembly zone in *syp-2*/+ oocytes, but these mutants did maintain full length SC after this zone (Figure 4C,E; P<0.001, Bonferroni P adjusted, Mann-Whitney). In contrast, *syp-2*/+ and *syp-3*/+ spermatocytes did not display any significant defects in SC assembly in early pachytene (Figure 4B,D). Additionally, *syp-2*/+ or *syp-3*/+ mutants in both sexes did not display any changes in SC disassembly or pachytene length, as indicated by DAPI morphology (Figure 4,S3). Thus, SC assembly and/or maintenance only in oocytes is sensitive to SYP-2 dosage during the early stages of pachytene.

**Figure 4:**
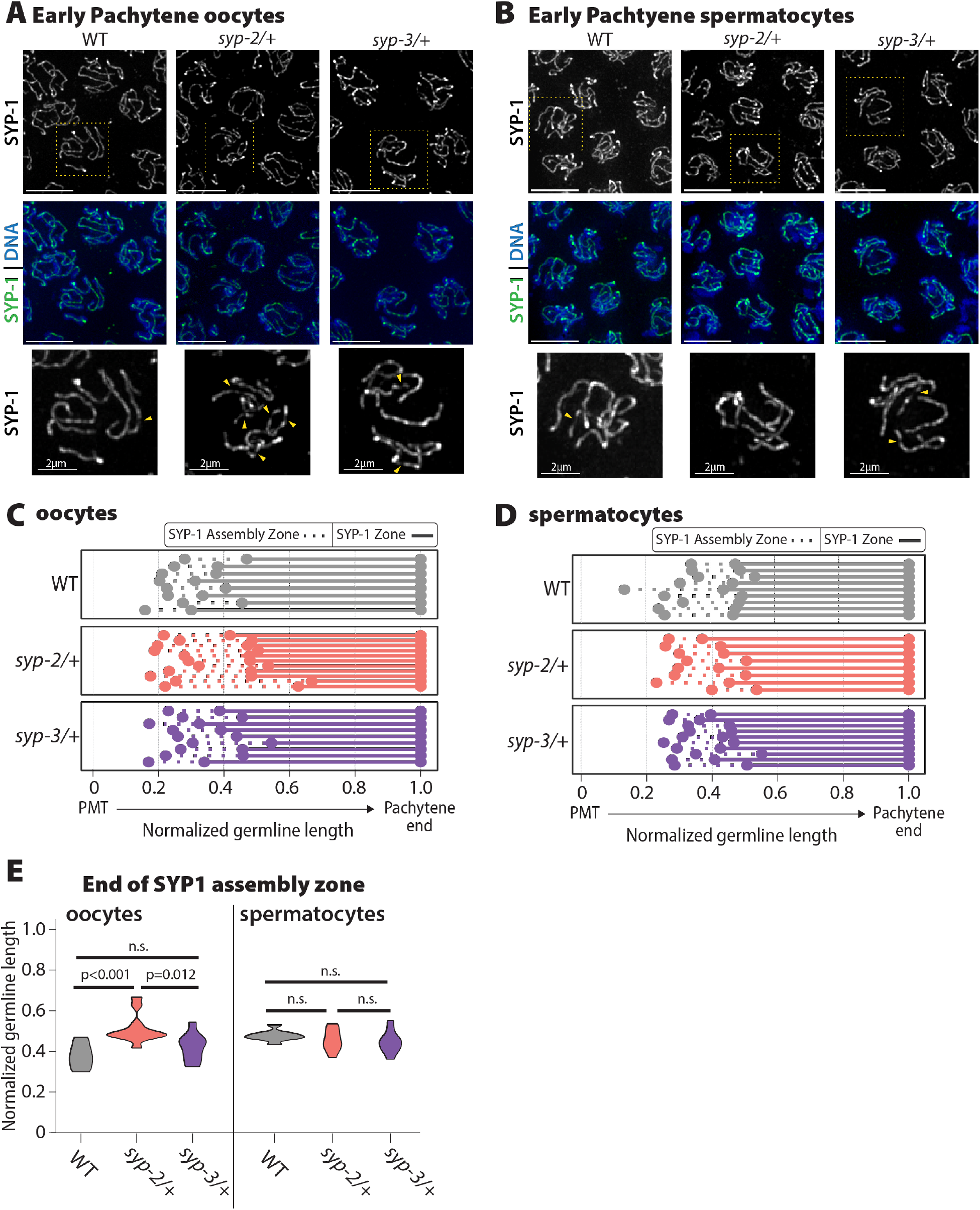
Oocyte SYP-1 assembly is uniquely sensitive to SYP-2 dosage. **(A**,**B)** Representative images early pachytene stained for SYP-1 in oocytes **(A)** and spermatocytes **(B)** from wild type (WT), *syp-2*/+, and *syp-3*/+. Yellow dashed box shows the nucleus that is enlarged below each merge image. Yellow arrowheads indicate regions where the SYP-1 signal is broken/not continuous. Scale bar represents 5µm **(C, D)** Measurement of the relative length of the SYP-1 assembly zone in oocytes **(C)** and spermatocytes **(D)** from the pre-meiotic tip (PMT) to the end of pachytene in wild type (WT, gray), *syp-2*/+ (pink), and *syp-3*/+ (purple). Dashed lines represent the SC assembly zone and solid lines represent fully assembled SC. **(E)** Quantification of the end of the SYP1 assembly zone in wild type (WT, gray), *syp-2*/+ (pink), and *syp-3*/+ (purple) in oocytes (left) and spermatocytes (right). All statistics are Bonferroni adjusted P values from Mann-Whitney U tests (n.s. = not significant). Oocyte data is from 8 wild type germlines, 10 *syp-2*/+ germlines, and 9 *syp-3*/+ germlines. Spermatocyte data is from 9 wild type germlines, 8 *syp-2*/+ germlines, and 10 *syp-3*/+ germlines.

### SYP-2 and SYP-3 dosage regulate recombination via separate, sex-specific mechanisms

Since altering SYP dosage permitted assembly of full length SC and did not impair chromosome pairing, we next assessed how reducing the dosage of SYP-2 and SYP-3 influenced recombination. To assess the mechanics of recombination, we used immunofluorescence to quantify and detect specific proteins that mark sites undergoing three different stages of recombination: (1) DSB formation with RAD-51 (Colaiacovo *et al*. 2003); (2) joint molecule formation with GFP::MSH-5 (Janisiw *et al*. 2018); and, (3) crossover designation with GFP::COSA-1 in oocytes (Yokoo *et al*. 2012) and OLLAS::COSA-1 in spermatocytes (Janisiw *et al*. 2018) (see Methods). For simplicity, here we refer to the tagged versions of GFP::MSH-5 and GFP::COSA-1 or OLLAS::COSA-1 as only the protein name either MSH-5 or COSA-1. The average number of foci of each protein was determined by using a sliding window of 0.01 along the normalized germline length, which was divided into early, mid, and late pachytene (see Methods).

#### RAD-51 foci are relatively unaffected by SYP dosage

Similar to previous studies, we found that the number and timing of RAD-51 foci per nucleus during pachytene is very different between the sexes (Figure 5A, 5B, S4, Table S2) (Jaramillo-Lambert and Engebrecht 2010; Checchi *et al*. 2014). Oocytes initiate DSBs later in the germline than spermatocytes, however, both sexes progressively repair these breaks throughout pachytene (Colaiacovo *et al*. 2003; Jaramillo-Lambert and Engebrecht 2010; Checchi *et al*. 2014; Toraason *et al*. 2021). Based on the RAD-51 foci counts, altering SYP dosage did not appear to have large effects on DSB initiation or subsequent progression through a repair pathway in either sex (Figure 5A, 5B, S4, Table S2). We did note changes in the number of RAD-51 foci specifically in oocytes of *syp-2*/+ and *syp-3*/+ during early pachytene (P<0.001; Figure 5A), but these slight alternations did not change the DSB repair dynamics with foci numbers declining at similar rates to wild type during pachytene progression. Additionally, we checked DSB-2 staining in oocytes, which marks the region of the germline where DSBs are induced and found no significant changes in the DSB-2 zone in either mutant (Figure S5). Thus, DSB formation and repair is not largely impacted by altering the dosage of SYP-2 or SYP-3.

**Figure 5:**
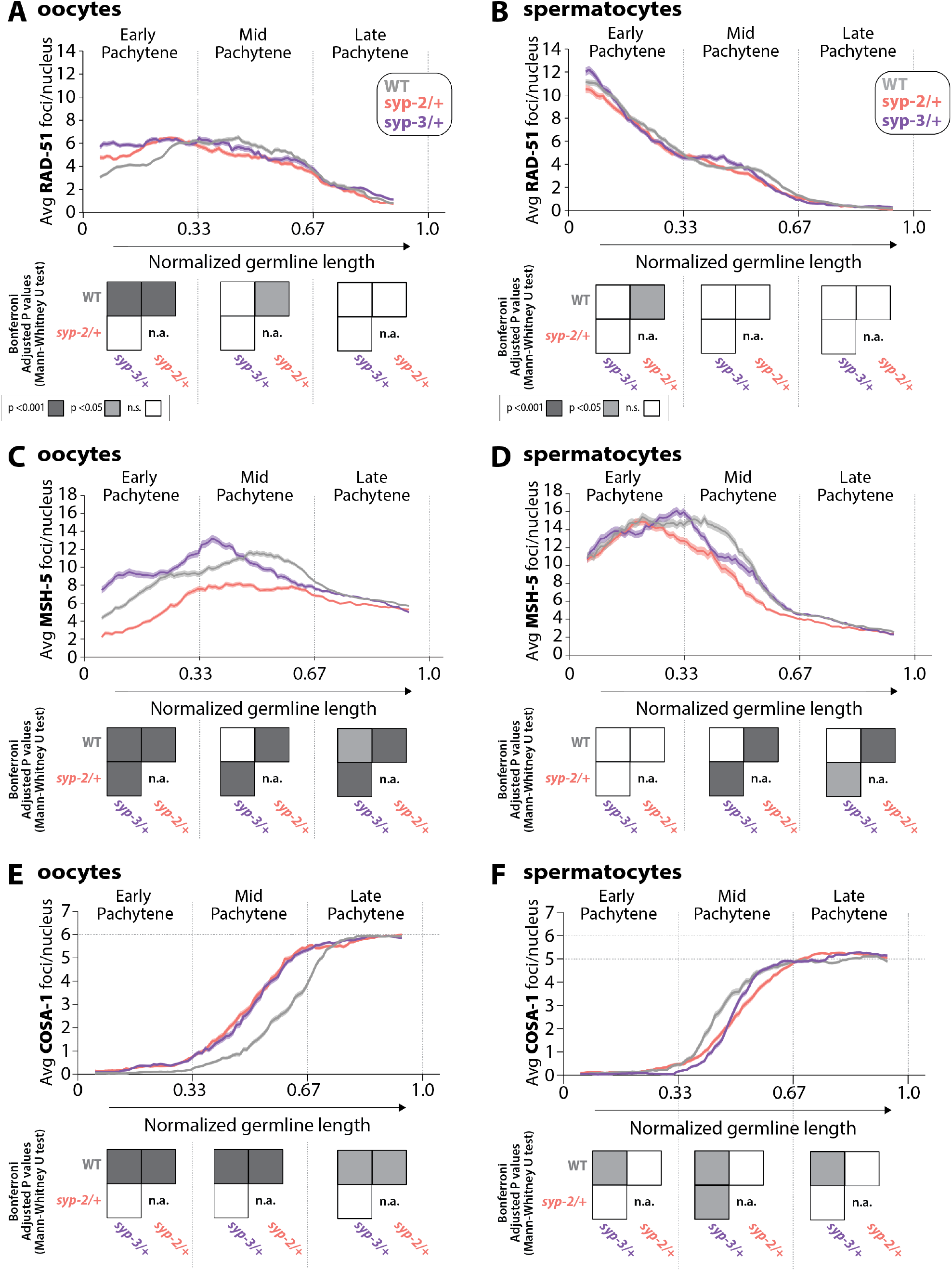
Sex-specific regulation of recombination by SYP-2 and SYP-3 dosage. **(A**,**B)** Quantification of the average number of RAD-51 foci per nucleus throughout pachytene in wild type (WT, gray), *syp-2*/+ (pink), and *syp-3*/+ (purple) from oocytes (**A**) and spermatocytes (**B**). **(C**,**D)** Quantification of the average number of MSH-5 foci per nucleus throughout pachytene in wild type (WT, gray), *syp-2*/+ (pink), and *syp-3*/+ (purple) from oocytes (**C**) and spermatocytes (**D**). **(E**,**F)** Quantification of the average number of COSA-1 foci per nucleus throughout pachytene in wild type (WT, gray), *syp-2*/+ (pink), and *syp-3*/+ (purple) from oocytes (**E**) and spermatocytes (**F**). Heat maps below each pachytene region show the Bonferroni adjusted P values from Mann-Whitney U tests with dark gray indicating p<0.001, light gray indicating p<0.05, and white indicating not significant (n.s.). The self-comparison between *syp-2*/+ was not determined (n.a.). n values for number of germlines and nuclei can be found in Table S2.

#### SYP dosage influences MSH-5 foci via sex-specific mechanisms

Consistent with the early loading of RAD-51 in spermatocytes, we found that MSH-5 is also loaded more quickly in spermatocytes than oocytes during pachytene (Figure 5C, 5D, S4, Table S2). Wild type spermatocytes reach peak amounts of MSH-5 foci around the transition between early and mid pachytene, whereas oocytes have peak amounts of MSH-5 foci in mid pachytene. Moreover, the peak amount of MSH-5 foci loaded per nucleus is higher in spermatocytes than oocytes (mean MSH-5 oocytes 11.6 foci ± SD 6.0 vs. spermatocytes 15.4 ± SD 7.0). These differences in MSH-5 foci between the sexes contribute to a growing body of work illustrating that the processing of recombination events is sexually dimorphic (Checchi *et al*. 2014; Brick *et al*. 2018; Li *et al*. 2020).

Reducing the dosage of SYP-2 and SYP-3 caused significant changes in the number of MSH-5 foci per nucleus during pachytene (Figure 5C, 5D, S4, Table S2), particularly in oocytes. *syp-2*/+ oocytes showed significant reductions in the average number of MSH-5 foci throughout pachytene, indicating that the amount of SYP-2 is crucial to load and/or maintain MSH-5 at a DSB (Figure 5C; P<0.001, Bonferroni adjusted P, Mann-Whitney). *syp-3*/+ oocytes displayed significant increases in the average number of MSH-5 foci per nucleus during early and mid pachytene (Figure 6C, P<0.001, Bonferroni adjusted P, Mann-Whitney). However, during mid pachytene, *syp-3*/+ oocytes rapidly lost MSH-5 foci earlier than wild type. Taken together, our data suggests that SYP-3 dosage regulates the timing of MSH-5 loading and off-loading (or maintenance at a DSB) in oocytes.

**Figure 6:**
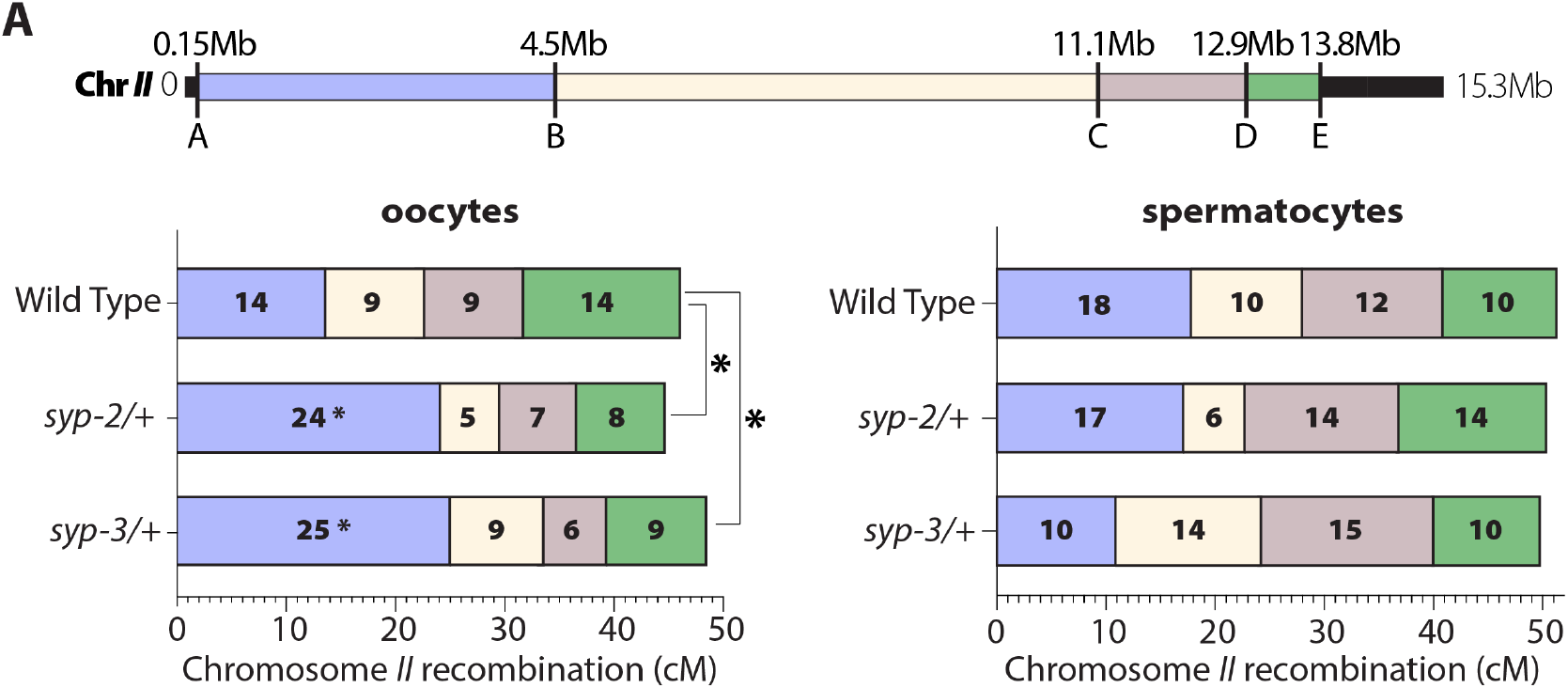
SYP dosage influences the crossover landscape in only oocytes. Recombination SNP mapping of Chromosome *II* in WT, *syp-2*/+, and *syp-3*/+ from oocytes (left) and spermatocytes (right) (see Methods for details). A diagram of the 15.3Mb Chromosome *II* shows the megabase location of each SNP assayed (A-E) and the colored boxed between each SNP show the intervals where crossovers were assessed. The map length (cM) is indicated in each crossover interval. The asterisks next to the map lengths indicate significance based on Fisher’s Exact tests compared between the mutants and wild type (*syp2*/+ P=0.0449; *syp3*/+ P=0.0170). The asterisks outside the bars indicate significance based on Chi-Squared tests between mutants and wild type (*syp2*/+ P=0.0343; *syp3*/+ P=0.0333). The worm counts for these plots can be found in Table 1.

In spermatocytes, only reducing the dosage of SYP-2 caused significant changes in MSH-5 foci during pachytene (Figure 5D, S4, Table S2). *syp-2*/+ spermatocytes initially form MSH-5 foci to levels similar to wild type in early pachytene. However, during mid and late pachytene *syp-2*/+ spermatocytes rapidly lose MSH-5 foci, suggesting that SYP dosage is important for the maintenance of MSH-5 at joint molecules (Figure 5D; P<0.001, Bonferroni adjusted P, Mann-Whitney). Notably, for both sexes the dosage of SYP-2 appears to be important for MSH-5 stability, suggesting a conserved role for SYP-2 in both sexes. Additionally, unlike oocytes, SYP-3 dosage does not change MSH-5 foci during pachytene in spermatocytes illustrating a sex-specific role for SYP-3 in regulating MSH-5 at DSB sites during recombination.

#### SYP dosage alters the timing of COSA-1 foci loading during pachytene

Unlike the sexually dimorphic DNA repair dynamics observed in RAD-51 and MSH-5, both oocytes and spermatocytes load COSA-1 in mid pachytene, and by late pachytene, all 6 COSA-1 foci in oocytes and 5 COSA-1 foci in spermatocytes have been established (Figure 5E, 5F, S4, Table S2). Unexpectedly, we noted that in wild type spermatocytes, a fraction of the nuclei in late pachytene also contained 6 or more COSA-1 foci (Figure S6). This was likely not observed in previous studies examining COSA-1 foci in males because the GFP::COSA-1 transgene used in those studies is driven by the *pie-1* promotor and the *cosa-1* 3’ UTR (Yokoo *et al*. 2012). Since *pie-1* promotors require specific 3’ UTRs, such as the *tbb-2* 3’ UTR, for robust spermatocyte expression (Merritt *et al*. 2008; Merritt *et al*. 2010), this GFP::COSA-1 transgene is only weakly expressed in males. Male worms do occasionally form double crossover events and indeed the nuclei with 6 or more COSA-1 foci appear to be double crossovers with two COSA-1 foci on one chromosome (Figure S6B) (Lim *et al*. 2008; Gabdank and Fire 2014). Our data demonstrate that double crossover events occur about ∼20% of the time in spermatocytes (Figure S6). Similar to previous studies, we also do not observe COSA-1 foci on the hemizygous X chromosome, which does not normally assemble the central region SYPs (Jaramillo-Lambert and Engebrecht 2010). Thus, the sex-specific changes we observed in SYP dynamics and accumulation may be influencing the sex-specific regulation of crossover number between oocytes and spermatocytes (Figures 1, 2, 3; see Discussion).

Altering the dosage of SYP-2 and SYP-3 cause significant changes in the timing of COSA-1 loading in both oocytes and spermatocytes (Figure 5E, 5F, S4, Table S2). In both *syp-2*/+ and *syp-3*/+ oocytes, the loading of COSA-1 foci was shifted to occur earlier in mid pachytene than wild type (Figure 5E; *syp-2*/+ P<0.001, *syp-3*/+ P<0.001, Bonferroni adjusted P, Mann-Whitney). In contrast, *syp-3*/+ spermatocytes exhibit a delay in the loading of COSA-1 foci during mid pachytene (Figure 5F; P<0.05, Bonferroni adjusted P, Mann-Whitney; Figure S7). Although *syp-2*/+ spermatocytes also displayed a potential delay in the loading of COSA-1, the difference is not statistically different from wild type (Figure 5F; P=1, Bonferroni adjusted P, Mann-Whitney). Taken together, dosage of SYP-2 and SYP-3 regulates the sex-specific timing of crossover designation.

By late pachytene, the final number of COSA-1 foci in oocytes (6 foci) and spermatocytes (5 or more foci) is not changed from the required one crossover per chromosome, thereby suggesting that SYP dosage does not largely influence the ability of each sex to designate the crossovers on all chromosomes (Figure 5E, 5F, S4, Table S2). Further, in oocytes, we observed 6 DAPI staining bodies at diakinesis for both *syp-2*/+ (30/30 oocytes with 6 DAPI bodies) and *syp-3*/+ (30/30 oocytes with 6 DAPI bodies) indicating that the 6 COSA-1 foci per nucleus at late pachytene mature into 6 crossovers that link the homologous chromosomes together at diakinesis.

### SYP-dosage regulates crossover landscape

Since we found that manipulating the dosage of SYP-2 and SYP-3 altered multiple steps of recombination and previous studies indicate that the SC can regulate crossover numbers (MacQueen *et al*. 2002; Colaiacovo *et al*. 2003; Smolikov *et al*. 2007b; Hayashi *et al*. 2010; Libuda *et al*. 2013; Gordon *et al*. 2021), we wanted to determine whether SYP-2 and SYP-3 dosage influences the recombination landscape by changing where crossovers are positioned along the length of the chromosome. To assess the recombination landscape, we used established single nucleotide polymorphism (SNP) recombination mapping between to two *C. elegans* isolates (Bristol and Hawaiian) to identify crossovers (see Methods) (Bazan and Hillers 2011). For oocytes, we mapped recombination on both Chromosome *II* and the *X* Chromosome. Since male worms have an unpaired *X* chromosome and do not form crossovers on this sex chromosome, only recombination on Chromosome *II* was mapped in spermatocytes.

On Chromosome *II*, both *syp-2*/+ and *syp-3*/+ altered the crossover landscape differently in each sex (Figure 6, Table 1). In spermatocytes, both *syp-2*/+ and *syp-3*/+ displayed no significant changes in crossover frequencies across all of Chromosome *II*. In contrast, both *syp-2*/+ and *syp-3*/+ oocytes displayed significant changes in the crossing over distribution on Chromosome *II* (Figure 6; *syp2*/+ P=0.0343, *syp3*/+ P=0.0333, Chi-Squared). Specifically, both mutants increased crossing over in the first interval (A to B) on the left side of Chromosome *II* by ∼10cM each (Figure 6; wild type 14cM, *syp-2*/+ 24cM, *syp-3*/+ 25cM; *syp2*/+ P=0.0449, *syp3*/+ P=0.0170, Fisher’s Exact). Intriguingly, this first interval is also where the pairing center is located on Chromosome *II*. Thus, one possible explanation for the elevated crossing over in the interval is that by reducing the SYP dosage, crossovers are now more often formed where the SC is first assembled at the pairing center (Hayashi *et al*. 2010; Rog and Dernburg 2015).

**Table 1:**
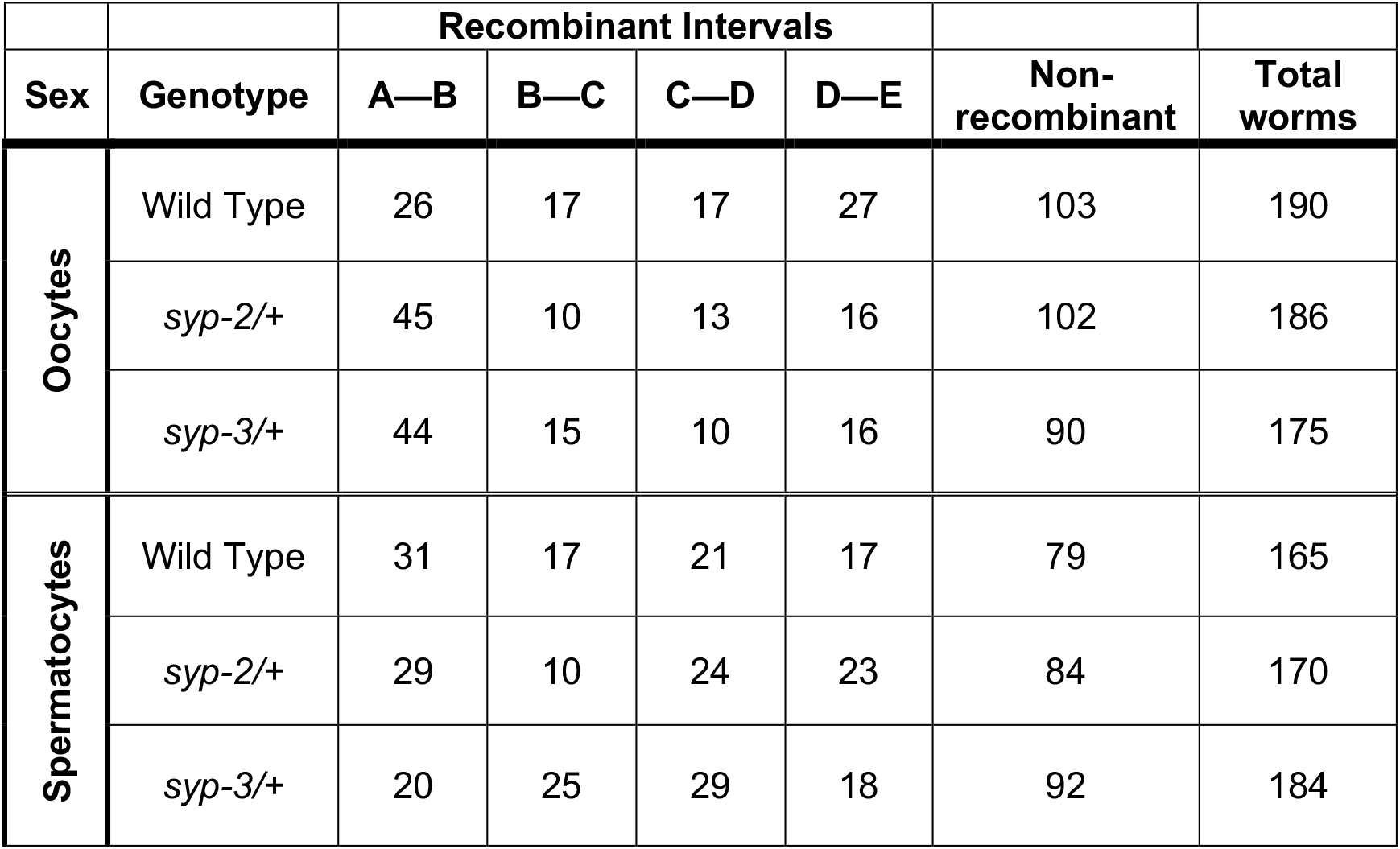
Chromosome *II* SNP mapping recombination.

On the *X* Chromosome, *syp-2*/+ oocytes display a significant decrease in crossover frequency along the entire chromosome (Figure S7A; Table S3; wild type 51cM, *syp-2*/+ 32cM; P=0.0343, Chi-Squared). This decrease was not observed in *syp-3*/+ oocytes, which showed no significant changes in crossover frequency to wild type for *X* Chromosome recombination (Figure S7A; Table S3; wild type 51cM, *syp-3*/+ 45cM; P=0.5849, Chi-Squared). The large decrease in crossing over in *syp-2*/+ suggests that these mutants should have a significant amount of *X* Chromosome nondisjunction or missegregation. In *C. elegans*, mutants with frequent *X* Chromosome nondisjunction produce a high incidence of male (Him) phenotype as male worms are hemizygous for the *X* Chromosome. Notably, *syp-2*/+ did not display an elevation in the frequency of male progeny nor did they have an increase in dead eggs (Figure S7B; wild type 0.4% dead eggs and 0.2% Him, *syp-2*/+ 0.4% dead eggs and 0.2% Him). Since the SNP recombination mapping experiment is performed in a hybrid Bristol/Hawaiian background, we also checked the hybrid background for male progeny and dead eggs. Due to known meiotic drive elements between these strains, the Bristol/Hawaiian hybrids produce more dead eggs (Seidel *et al*. 2008; Seidel *et al*. 2011). However, *syp-2*/+ Bristol/Hawaiian hybrids did not display a higher incidence of dead eggs or more male progeny than the wild type Bristol/Hawaiian hybrid (Figure S7B; wild type 19.4% dead eggs and 0.2% Him; *syp-2*/+ 23.3% dead eggs and 0.1% Him). Thus, it remains unclear as to why reducing SYP-2 causes a significant decrease in recombination without a corresponding increase in *X* chromosome nondisjunction. One possible explanation is that *syp-2*/+ have a chromosome distortion event occurring in the later stages of meiosis that is removing recombinant *X* chromosomes. Future studies are needed to understand what is happening to the recombinant *X* chromosomes in *syp-2*/+.

In comparison to wild type, the broods from *syp-3*/+ hermaphrodites exhibited both an increased lethality (more dead eggs; Figure S7B, wild type Bristol 0.4% and hybrid 19.4% dead eggs; *syp-3*/+ Bristol 1.2% and hybrid 27.0% dead eggs) as well as an higher incidence of male progeny than wild type (Figure S7B, wild type Bristol 0.2% and hybrid 0.2% HIM; *syp-3*/+ Bristol 0.7% and hybrid 1.6% HIM, Bristol p<0.0001, Hybrid p<0.0001, Chi-Squared). In addition, *syp-3*/+ also produced a higher rate of progeny with dumpy and/or uncoordinated mutant phenotypes than both wild type and *syp-2*/+ (Figure S7B, Bristol 0.06% mutant progeny, *syp-2*/+ Bristol 0.06% mutant progeny, *syp-3*/+ Bristol 0.47% mutant progeny; wild type Bristol p=0.0207, *syp-2*/+ Bristol p=0.0388, Fisher’s Exact). SYP proteins have been implicated in regulating DSB repair pathway choice by preventing access to error prone and sister chromatid repair (Smolikov *et al*. 2007a; Rosu *et al*. 2011; Lemmens *et al*. 2013; Yin and Smolikove 2013; Macaisne *et al*. 2018; Lascarez-Lagunas *et al*. 2022). Additionally, SYP-3 directly interacts with the pro-crossover protein BRC-1 to promote crossover DSB repair (Janisiw *et al*. 2018). Thus, the mutant progeny observed from *syp-3*/+ hermaphrodites are likely due to error prone repair using nonhomologous end joining, which is used by *syp-3* nulls to repair persisting DSBs (Smolikov *et al*. 2007b). These results suggest that SYP-3 amounts are important for suppressing error prone repair.

### SYP-2 and SYP-3 dosage influences the SC composition during pachytene

The SYP dosage-dependent regulation of recombination (Figures 4, 5), different patterns of SYP protein incorporation throughout pachytene (Figure 2), and independent regulation of SYP loading into the SC (Figures 2, 3) suggest that SYP accumulation in the SC may be dynamically altered during pachytene to regulate the steps of recombination. Thus, reducing the dosage of each SYP may influence how the SYPs are accumulating within the SC, which could influence the regulation of recombination. To determine how dosage of each SYP influences the incorporation of each SYP during pachytene, we used heterozygous null mutants of each *syp* with a wild type fluorescently tagged GFP::SYP-2 or mCherry::SYP-3: *syp-2(ok307)/gfp::syp-2* (referred to as *syp-2*/+) or *syp-3(ok785)/mCherry::syp-3* (referred to as *syp-3*/+). Indeed, the dosage of SYP-2 and SYP-3 influenced both SYP-2 and SYP-3 accumulation during pachytene in both sexes (Figure 7, Table S1).

**Figure 7:**
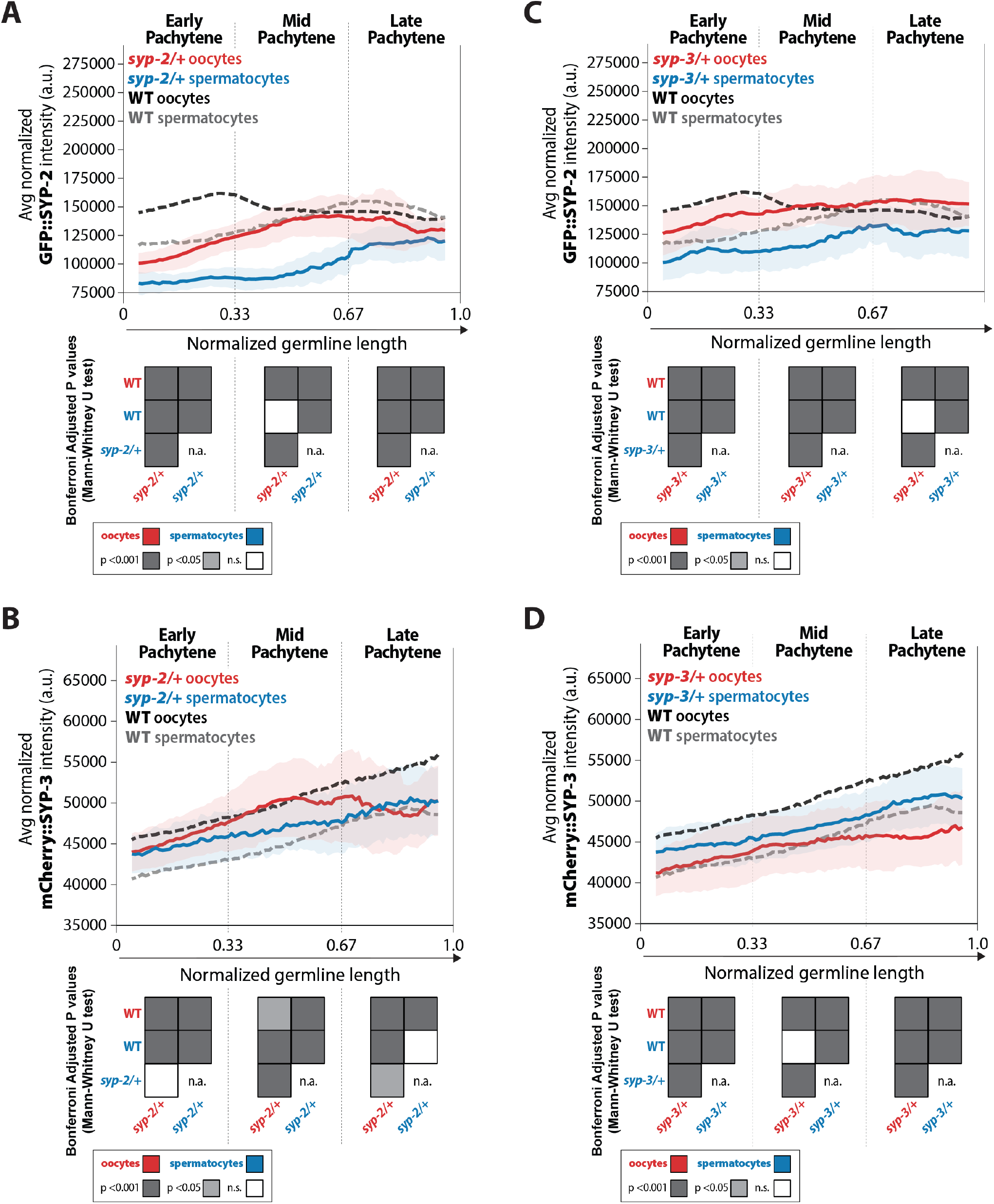
Dosage of SYP-2 and SYP-3 influence the amount of each SYP within the SC via sex-specific mechanisms. **(A**,**B)** Quantification of the mean intensity of GFP::SYP-2 **(A)** and mCherry::SYP-3 **(B)** per nucleus normalized by the volume of each nucleus (see Methods) throughout pachytene for *syp-2*/+. **(C**,**D)** Quantification of the mean intensity of GFP::SYP-2 **(C)** mCherry::SYP-3 **(D)** per nucleus normalized by the volume of each nucleus throughout pachytene (normalized, see Methods) for *syp-3*/+. Oocytes are shown in red with the standard deviation shown as a pale red band and spermatocytes are shown in blue with the standard deviation shown as a pale blue band. The mean intensity of GFP::SYP-2 **(A**,**B)** and mCherry::SYP-3 **(C**,**D)** for wild type (WT) oocytes (black) and wild type (WT) spermatocytes (gray) are shown as dashed lines. Heat maps below each pachytene region show the Bonferroni adjusted P values from Mann-Whitney U tests with dark gray indicating p<0.001, light gray indicating p<0.05, and white indicating not significant (n.s.). The self-comparison between spermatocyte *syp-2*/+ or *syp-3*/+ mutants was not determined (n.a.). n values for number of germlines and nuclei can be found in Table S1.

Altering the dosage of SYP-2 caused both sexes to initially have a ∼0.7-fold decrease in SYP-2 amounts within the SC, however, SYP-2 levels progressively increased throughout pachytene until wild type amounts were reached or pachytene ended (Figure 7A, late pachytene oocytes 0.94-fold change from wild type; spermatocytes 0.80-fold change from wild type). This result suggests that both sexes attempt to compensate for the haploinsufficiency of losing a *syp-2* gene by loading as much SYP-2 as possible during pachytene. In contrast, SYP-3 accumulation in *syp-2*/+ mutants displayed sex-specific responses (Figure 7C). In oocytes, reducing the SYP-2 dosage caused a slight decrease in SYP-3 levels in early pachytene (0.97-fold change from wild type) that becomes more pronounced in late pachytene (0.91-fold change from wild type), which we noted was the same window when SYP-2 levels reached wild type amounts in *syp-2*/+ mutants (Figure 7C; P<0.001, Bonferroni adjusted, Mann-Whitney). In spermatocytes, reducing the dosage of SYP-2 caused an increase in SYP-3 amount in early pachytene (1.1-fold change from wild type) that are indistinguishable from wild type by late pachytene (1.0-fold change from wild type) (Figure 7C). Thus, while altering SYP-2 dosage caused both sexes to respond similarly with reduced loading of SYP-2 into assembled SC, SYP-3 levels within assembled SC were inversely affected via sex-specific mechanisms.

Notably, altering SYP-3 dosage caused the same overall changes to SYP-2 and SYP-3 amounts in each sex as altering SYP-2 dosage, albeit the degree in which SYP changed was different (Figure 7B, 7D, Table S1). Specifically, reducing SYP-3 dosage caused an initial reduction of SYP-2 amounts in both sexes that was less than the reduction in *syp-2*/+ (Figure 7A, 7B; early pachytene *syp-2*/+ oocytes 0.72-fold change vs. *syp-3*/+ oocytes 0.87-fold change; early pachytene *syp-2*/+ spermatocytes 0.71-fold change vs. *syp-3*/+ spermatocytes 0.89-fold change). However, similar to *syp-2*/+, both sexes progressively increase SYP-2 amounts in *syp-3*/+ until wild type levels were reached or pachytene ended (Figure 7A, 7B, late pachytene *syp-2*/+ oocytes 0.94-fold change vs. *syp-3*/+ oocytes 1.1-fold change; late pachytene *syp-2*/+ spermatocytes 0.80-fold change vs. *syp-3*/+ spermatocytes 0.86-fold change). In oocytes, reducing SYP-3 dosage caused a stronger decrease in SYP-3 amounts compared to *syp-2*/+ (Figure 7C, 7D, early pachytene *syp-2*/+ 0.97-fold change vs *syp-3*/+ 0.91-fold change, mid pachytene 0.99-fold change vs *syp-3*/+ 0.90-fold change, late pachytene 0.91-fold change vs. *syp-3*/+ 0.85-fold change). In contrast, *syp-3*/+ spermatocytes displayed an increase in SYP-3 amounts that was largely similar to the increase in SYP-3 observed in *syp-2*/+ spermatocytes (Figure 7C, 7D, early pachytene *syp-2*/+ 1.1-fold change vs *syp-3*/+ 1.1-fold change, mid pachytene 1.0-fold change vs *syp-3*/+ 1.0-fold change, late pachytene 1.0-fold change vs. *syp-3*/+ 1.0-fold change).

Taken together, the dosage of each SYP influences the accumulation of SYP-2 and SYP-3 within the assembled SC similarly for each sex. Oocytes respond to reduced SYP dosage by decreasing both SYP-2 and SYP-3 in the SC, while spermatocytes respond to SYP reductions by decreasing SYP-2 and increasing SYP-3 in the SC. These differences are likely influencing the changes we observed in recombination for both *syp-2*/+ and *syp-3*/+ (Figure 5) and suggest that the dosage of each SYP throughout pachytene regulates recombination in a sex-specific manner (see Discussion, Figure 8).

**Figure 8:**
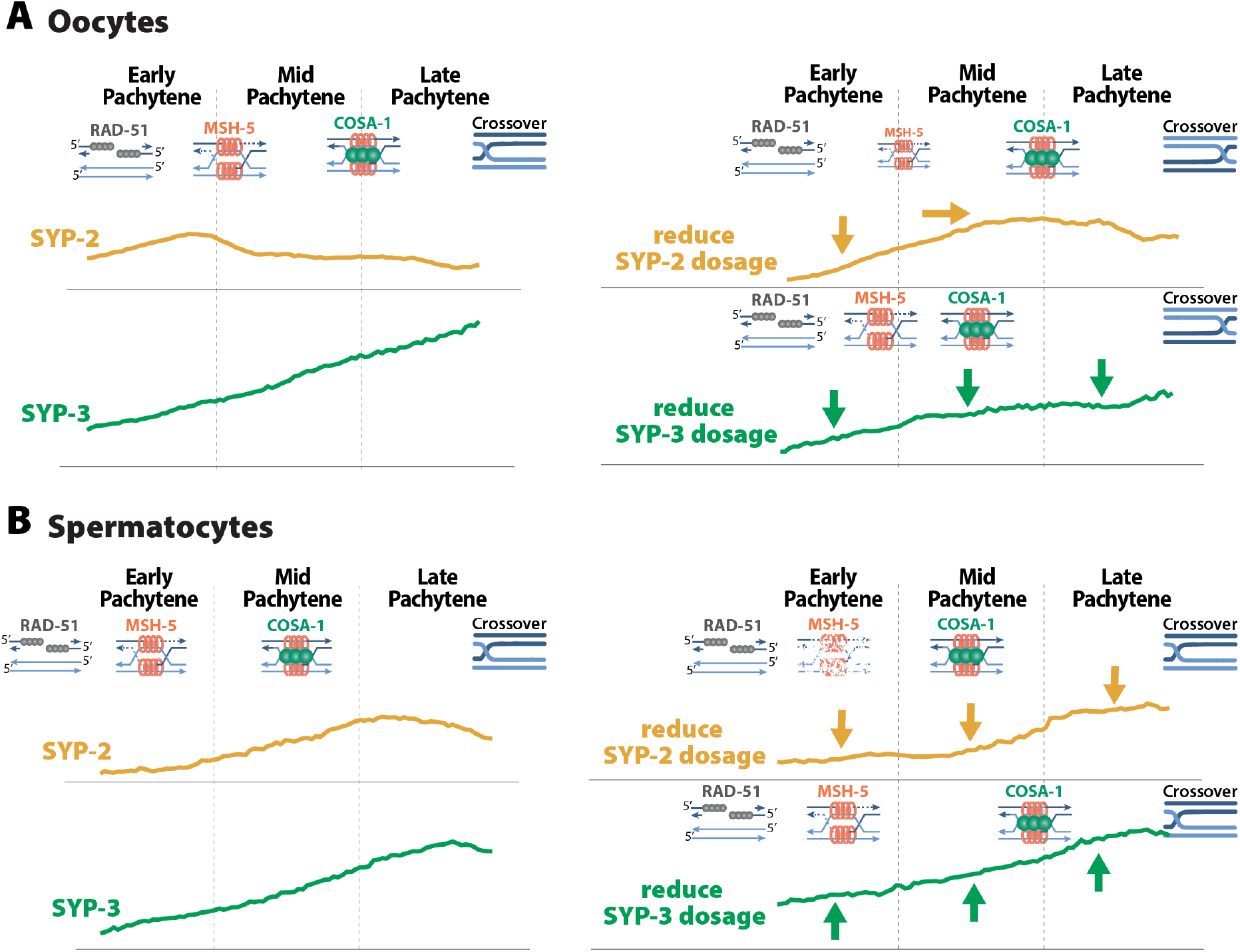
SYP dosage influences the sexually dimorphic regulation of recombination. **(A)** In oocytes, SYP-2 amounts with the SC are critical for the proper formation and/or maintenance of joint molecules stabilized by MSH-5. Reducing the amount of SYP-2 dosage causes decreases in the amount of SYP-2 in early pachytene and shifts the peak amounts of SYP-2 toward mid/late pachytene. This alteration to SYP-2 composition with the SC causes severe reduction in MSH-5. SYP-3 amounts in the SC are important for the proper timing of recombination. When the dosage of SYP-3 is reduced, SYP-3 composition in the SC is reduced throughout pachytene causing faster resolution of jointed molecules and faster designation of crossovers during pachytene. This ultimately causes changes in the recombination landscape where crossovers are more often position near the pairing center. **(B)** In spermatocytes, SYP-2 amounts are important for the maintenance of MSH-5 stabilized joint molecules. When SYP-2 dosage is reduced, the amount of SYP-2 in the SC is reduced and MSH-5 foci are rapidly lost either because they are resolved quickly or the stability of the joint molecules is compromised. SYP-3 amounts in spermatocytes also influence the timing of recombination but when SYP-3 dosage is reduced SYP-3 amounts are increased rather than decreased. Thus, elevated SYP-3 levels in spermatocytes cause a delay in crossover designation in spermatocytes.

## DISCUSSION

### Sexually dimorphic regulation of recombination by the SC

One of the roles for the SC during meiosis is in regulating recombination to promote the establishment of a crossover on each homolog pair. Our data here demonstrate that the SC in *C. elegans* regulates recombination via sexually dimorphic mechanisms. We show that the central region proteins SYP-2 and SYP-3 have sex-specific differences in protein turnover rates within the SC and the incorporation of SYP proteins within the SC differ between oocytes and spermatocytes. Moreover, we find that these sex-specific differences in SYP-2 and SYP-3 amounts in the SC are important for the regulation of recombination. Specifically, our data support a model where the amount of SYP-2 and SYP-3 within the SC regulates the proper timing and execution of specific steps of recombination (Figure 8).

Our data suggest that the amount of SYP-2 within the SC of oocytes promotes the formation and/or maintenance of joint molecules (Figure 8A). *syp-2*/+ oocytes incorporate less SYP-2 in the SC during meiotic stages when MSH-5 stabilized joint molecules are formed (Figures 5, 7) and also exhibit a severe reduction in MSH-5 foci (Figure 5). This relationship between the SC and joint molecules is SYP-2 specific, as a reduction in SYP-3 levels within an assembled SC did not result in reduced MSH-5 foci (Figures 5, 7). We therefore suggest that a specific threshold of SYP-2 in the SC ensures the formation and/or stabilization of joint molecules.

Similar to oocytes, spermatocytes SYP-2 dosage is required for the maintenance of joint molecules marked by MSH-5 (Figure 8B). Unlike oocytes, reducing SYP-2 levels in spermatocytes did not alter the assembly of the SC (Figure 4) and is likely why MSH-5 foci are able to initially form near wild type levels in early pachytene (Figure 5). However, the rapid loss of MSH-5 in spermatocytes during mid pachytene suggests that these joint molecules are either being rapidly resolved and designated for crossover recombination or destabilized, resulting in MSH-5 removal. We favor a model where the rapid loss of MSH-5 foci is underpinned by destabilization of joint molecules, as the crossover designation marker COSA-1 is not prematurely loaded when SYP-2 dosage is reduced. This function of SYP-2 in the maintenance of joint molecules appears conserved between the sexes, as *syp-2*/+ spermatocytes initially form wild type levels of MSH-5 foci that are rapidly lost (Figure 5). However, the specific threshold of SYP-2 required in spermatocytes to establish and/or stabilize joint molecules is significantly reduced compared to oocytes (Figures 2, 5, 7). This discrepancy between the sexes suggests that there are mechanistic differences in how joint molecules are regulated between spermatocytes and oocytes. Thus, our work here adds to a growing body of work illustrating that spermatocytes process DSBs into crossovers differently than oocytes (Jaramillo-Lambert *et al*. 2007; Jaramillo-Lambert and Engebrecht 2010; Checchi *et al*. 2014; Brick *et al*. 2018; Li *et al*. 2020).

We further propose that SYP-3 dosage in oocytes regulates the timing of recombination steps (Figure 8A). SYP-3 accumulates within the SC throughout pachytene as recombination intermediates are successively processed (Figure 2). Notably, MSH-5 marked joint molecules and COSA-1 marked crossover designation appear prematurely when the levels of SYP-3 in the SC are reduced (Figures 5, 7). This acceleration in crossover designation coincides with a disproportionate formation of crossovers near the pairing center side of Chromosome *II* (Figure 6). This change in crossover designation and positioning was also present in *syp-2*/+. However, we assert that this phenotype is likely underpinned by SYP-3 levels, as SYP-3 incorporation is also reduced in *syp-2*/+ (Figure 7). Thus, the timing of recombination events in the germline appear to be sensitive to the amounts of SYP-3. We suggest that SYP-3 incorporation is dynamically regulated in response to meiotic stresses to ensure that recombination is completed by the end of pachytene, which is spatially and temporally limited by the length of the gonad.

In spermatocytes, SYP-3 also regulates the timing of recombination events during pachytene. However, spermatocytes SYP-3 incorporation is minimally impacted by *syp-3*/+ heterozygosity (Figure 8B). We suggest that the relatively normal levels of SYP-3 in the *syp-3*/+ spermatocyte SC explains the absence of crossover positioning defects (Figure 7). In fact, SYP-3 levels are subtly elevated in early pachytene in *syp-3*/+ spermatocytes and coincides with a delay in crossover designation (Figures 5, 7). As spermatogenesis and oogenesis operate on very different timescales and the amount of SYP-3 in the SC of spermatocytes and oocytes differs, we raise the possibility that titrating the level of SYP-3 incorporation may function to regulate the timing of recombination events between the sexes.

### Spermatocytes regulate the SC differently than oocytes

Sex comparative studies are critical to understand the differences in egg and sperm development. Here we demonstrate that the SC in *C. elegans* is sexually dimorphic. Intriguingly, the sex-specific differences in SYP-2 and SYP-3 dynamics within the SC suggest a difference in protein regulation between the sexes (Figure 1). The progressive stabilization of SC during pachytene in oocytes has been linked with phosphorylation of specific SYP proteins (Nadarajan *et al*. 2017; Pattabiraman *et al*. 2017). Additionally, during oogenesis, many SYP proteins are known to be post-translationally modified, including SYP-2, in response to recombination and many of these modifications are critical for DSB formation and repair as well as the timing of SC assembly and disassembly (Nadarajan *et al*. 2016; Nadarajan *et al*. 2017; Sato-Carlton *et al*. 2018; Garcia-Muse *et al*. 2019; Lascarez-Lagunas *et al*. 2022). Since spermatocytes also displayed the same progressive stabilization of the SYP-2 and SYP-3 during pachytene as oocytes, it is possible that the same post-translational modifications of SYP proteins may also have similar functions in spermatocytes.

We found that SYP-2 was more dynamic in spermatocytes even at the “stabilized” state in late pachytene (Figure 1). Thus, spermatocytes may not stabilize the SC to the same degree as oocytes. One reason for this could be to allow DSBs in late pachytene to be repaired with the homolog. In oocytes, the stabilization of the SC in late pachytene is thought to prevent the formation of more crossovers with the homolog. Spermatocytes have been shown to have differences in DNA repair, the number of crossovers, and the checkpoints that monitor repair events, and these differences may require a more dynamic SC (Jaramillo-Lambert *et al*. 2007; Jaramillo-Lambert and Engebrecht 2010; Jaramillo-Lambert *et al*. 2010; Checchi *et al*. 2014; Gabdank and Fire 2014; Li *et al*. 2020). Future studies are needed in spermatocytes to examine these relationships between SC dynamics, post translation modification of the SC, and DNA repair outcomes.

The amount of SYP-2 and SYP-3 required to properly execute the same steps of recombination is significantly different between oocytes and spermatocytes (Figure 8). Oocytes require more of both SYP-2 and SYP-3 during early and mid pachytene than spermatocytes (Figures 2, 6). One possibility for this difference is due to male worms having a hemizygous X chromosome, which does not normally assemble the SC (Jaramillo-Lambert and Engebrecht 2010). This lack of SC on one chromosome should potentially reduce the amount of SC proteins we get from our analysis. If this were the case, then we would expect the SYP proteins to be reduced in spermatocytes along the entire length of pachytene. Instead, we SYP-2 levels in spermatocytes reach those of oocytes during late pachytene. Further, spermatocytes have increased SYP-2 turnover dynamics in late pachytene compared to oocyte SYP-2 (Figure 1), which may help facilitate the repair of DSBs in late pachytene since spermatocytes do not undergo checkpoint-mediated apoptosis (Jaramillo-Lambert *et al*. 2010). Thus, having a more dynamic SYP-2 may aid DSB repair when errors occur late in pachytene.

Our observation that reduced SYP-3 dosage triggers an increase in SYP-3 composition within the SC suggests that spermatocytes may be able to upregulate the expression of *syp-3* to compensate for the loss of a functional copy of *syp-3*. Given that spermatocytes differ in the dynamics of protein turnover within the assembled SC, one way spermatocytes could alter SYP-3 levels is to alter the SYP-3 turnover rates within the SC. We demonstrate that spermatocytes do indeed have altered protein dynamics in the SC compared to oocytes and that SYP-2 and SYP-3 have different dynamics within each sex (Figure 1). Therefore, differences in the dynamics both between the sexes and between SYP-2 and SYP-3 could be used to regulate the composition of the SC throughout pachytene. Future studies are necessary to understand these sex-specific differences examining how the SYP proteins are regulated in each sex both at the gene expression level as well as at the protein dynamics level.

### Independent regulation of SYP-2 and SYP-3

One common feature amongst SC proteins in nearly all SC-containing organisms is that most, if not all, of the central region proteins are dependent upon each other for assembly of the SC (reviewed in Cahoon and Hawley 2016). In *C. elegans*, the SYP proteins are also dependent on each other for protein stability. Specifically, the result where depletion of one SYP leads to the degredation of the other SYPs, has led to the assumption that the SYPs are all completely interdependent for assembly, stability and function (Colaiacovo *et al*. 2003). Here we show that fluctuations in both SYP-2 and SYP-3 levels can not only differentially influence the amount of each other within the SC (Figure 7), but also each SYP protein can be regulated independently (Figures 2, 3, 7). Notably, we show that these differences in the proportion of each SYP within the SC is directly involved in regulating specific steps of recombination. Thus, we hypothesize that each SYP in the SC maintains both SYP-dependent functions where they function together as a group (*e*.*g*. assembling the SC) and SYP-independent functions where they can individually influence different aspects of meiosis (*e*.*g*. regulating specific steps of recombination).

In worms and yeast, the SC appears to grow in width throughout pachytene suggesting that the composition of proteins within the complex is highly dynamic (Voelkel-Meiman *et al*. 2012; Pattabiraman *et al*. 2017). Here we found that the incorporation of SYP-2 and SYP-3 are not proportional during pachytene (Figure 1). Further, our results suggest that recombination influences the pattern of SYP accumulation within the SC and that each of the SYPs are independently regulated within the SC (Figure 3). The ability to adjust the amount of SYP-2 and SYP-3 within the SC demonstrates flexibility in the requirements of SYP-2 and SYP-3 to assemble the complex. SYP-2 is positioned in the very center of the SC and may be slightly more external on the complex to SYP-3 (KÖhler *et al*. 2020), which would make it easier to alter the amount of SYP-2 without compromising the whole complex. Also, the proteins within this very central region of the SC play a critical role in recombination (Voelkel-Meiman *et al*. 2015; Voelkel-Meiman *et al*. 2019; Gordon *et al*. 2021; Voelkel-Meiman *et al*. 2022). Thus, these proteins in the center of the SC, like SYP-2, may not have as strong of structural roles in the complex as those positioned broadly in the central region, like SYP-3.

Not only are the patterns of SYP-2 and SYP-3 accumulation in the SC different, but the dosage of each SYP protein can influence the amount of the other protein. Regardless of which SYP dosage was altered, the same change in SYP-2 and SYP-3 accumulation occurred individually for each sex. In oocytes, both SYP-2 and SYP-3 amounts decreased in response to reduced SYP-2 or SYP-3 (Figure 7). Whereas in spermatocytes, SYP-2 amounts decreased while SYP-3 amounts increased in response to reduced SYP-2 or SYP-3 (Figure 7). One explanation for these sex-specific differences is that spermatocytes require a different stoichiometry of proteins in the SC compared to oocytes. In mice, the SC in oocytes is narrower than the SC between spermatocytes due to structural differences in the organization of proteins within the central element and the chromosome axis (Agostinho *et al*. 2018). While *C. elegans* does not have a defined central element based on electron microscopy images of the SC, SYP-2 is located in the very center of the SC and this is where the central element proteins are located in other organisms (Cahoon and Hawley 2016). Future studies to determine the stoichiometric ratios of chromosome axis proteins may reveal that spermatocytes assemble an SC that is structurally different from oocytes (Woglar *et al*. 2020).

## ACKNOWLEDGEMENTS

We thank the CGC for strains, which is funded by NIH Office of Research Infrastructure Programs (P40 OD010440). We thank members of the Libuda Lab for discussion and comments on the manuscript and the Nicola Silva lab for the SYP-1 antibody. This work was supported by the National Institutes of Health R35GM128890 to DEL and a Jane Coffin Childs Postdoctoral Fellowship and National Institutes of Health 1K99HD109505-01 to CKC. DEL is also a Searle Scholar and recipient of a March of Dimes Basil O’Connor Starter Scholar award.

## METHODS

### C. elegans strains, genetics, CRISPR, and culture conditions

All strains were generated from the N2 background and were maintained and crossed at 20**°**C under standard conditions on nematode growth media (NGM) with lawns of *Escherichia coli* (*E. coli*). In Vivo Biosystems tagged the C-terminus of SYP-3 with a piRNA optimized mCherry using CRISPR/Cas9. The CRISPR homology-directed repair template was constructed containing at least 500 base pairs of homology on either side of the insertion site at the SYP-3 locus. A small region of DNA was recoded section at the sgRNA site to avoid Cas9 cutting the template and mCherry was attached to SYP-3 with a glycine serine linker (GGSGGGGS). These repair constructs were synthesized into plasmids and injected into the worm with two sgRNAs. All sequences and screening primers for the CRISPR/Cas9 tagging of SYP-3 are in Table S#. CRISPR/Cas9 worm lines were backcrossed to N2 worms three times before processing with any strain construction. The following strains were used in this study:

N2: Bristol wild type strain.

CB4856: Hawaiian wild type strain.

DLW114: *unc-18(knu969[unc-18::AID*]) X. reSi7 [rgef-1p::TIR1::F2A::mTagBFP2::AID*::NLS::tbb-2 3’UTR] I*.

DLW118: *unc-18(knu969[unc-18::AID*]) X. reSi7 [rgef-1p::TIR1::F2A::mTagBFP2::NLS::AID*::tbb-2 3’UTR] I. GFP::syp-2 V*.

DLW119: *syp-3(knu999[mCherry::syp-3]) II*.

DLW128: *unc-18(knu969[unc-18::AID*]) X. reSi7 [rgef-1p::TIR1::F2A::mTagBFP2::NLS::AID*::tbb-2 3’UTR] syp-3(knu999[mCherry::syp-3]) I*.

DLW160: *unc-18(knu969[unc-18::AID*]) X. reSi7 [rgef-1p::TIR1::F2A::mTagBFP2::AID*::NLS::tbb-2 3’UTR] syp-3(knu999[mCherry::syp-3]) I. cosa-1(tm3298)/sC1(s2023) [dpy-1(s2170) umnIs41] III. GFP::syp-2 V*.

DLW163: *unc-18(knu969[unc-18::AID*]) X. reSi7 [rgef-1p::TIR1::F2A::mTagBFP2::AID*::NLS::tbb-2 3’UTR]. spo-11(me44)/nT1 [qls51] IV. GFP::syp-2/nT1 [qls51] V*.

DLW188: *syp-2(ok307)/tmC16 [unc-60(tmIs1210)] V*.

DLW190: *syp-3(ok785)/tmC18 [dpy-5(tmIs1200)] I*.

DLW192: *unc-18(knu969[unc-18::AID*]) X. reSi7 [rgef-1p::TIR1::F2A::mTagBFP2::NLS::AID*::tbb-2 3’UTR] syp-3(knu999[mCherry::syp-3]) I. spo-11(me44)/nT1 [qls51] IV. GFP::syp-2/nT1 [qls51] V*.

DLW193: *unc-18(knu969[unc-18::AID*]) X. reSi7 [rgef-1p::TIR1::F2A::mTagBFP2::NLS::AID*::tbb-2 3’UTR] syp-3(knu999[mCherry::syp-3]) I. GFP::syp-2 V*.

DLW195: *meIs8[unc-119(+) pie-1promoter::gfp::cosa-1] II. syp-2(ok307)/tmC16 [unc-60(tmIs1210)] V*.

DLW196: *meIs8[unc-119(+) pie-1promoter::gfp::cosa-1] II. syp-3(ok785)/tmC18 [dpy-5(tmIs1200)] I*.

DLW197: *unc-18(knu969[unc-18::AID*]) X. reSi7 [rgef-1p::TIR1::F2A::mTagBFP2::NLS::AID*::tbb-2 3’UTR] syp-3(knu999[mCherry::syp-3]) I. syp-2(ok307)/tmC16 [unc-60(tmIs1210)] V*.

DLW198: *unc-18(knu969[unc-18::AID*]) X. reSi7 [rgef-1p::TIR1::F2A::mTagBFP2::NLS::AID*::tbb-2 3’UTR] syp-3(ok785)/hT2 [bli-4(e937) let-?(q782) qIs48] (I;III). GFP::syp-2 V*.

DLW208: *msh-5[ddr22(GFP::msh-5)] IV; syp-2(ok307)/tmC16 [unc-60(tmIs1210)] V*.

DLW209: *syp-3(ok785)/tmC18 [dpy-5(tmIs1200)] I; msh-5[ddr22(GFP::msh-5)] IV*.

DLW211: *cosa-1[ddr12(OLLAS::cosa-1)] III; syp-2(ok307)/tmC16 [unc-60(tmIs1210)] V*.

DLW212: *syp-3(ok785)/tmC18 [dpy-5(tmIs1200)] I; cosa-1[ddr12(OLLAS::cosa-1)] III*.

AV630: *meIs8[unc-119(+) pie-1promoter::gfp::cosa-1] II*.

NSV97: *cosa-1[ddr12(OLLAS::cosa-1)] III*.

NSV129: *msh-5[ddr22(GFP::msh-5)] IV*.

### Microscopy

Worms were mounted for all live imaging studies using our auxin-inducible conditional paralysis method, which is described in (Cahoon and Libuda 2021). Briefly, young adult worms (18-24hrs post L4) from a parental generation that was grown on NGM with either 1mM auxin (for oocyte studies) or 10mM auxin (for spermatocyte studies) were picked into 1µL drop of live imaging media (M9 media with 25mM serotonin (Sigma-Aldrich, cat no. H7752, (Rog and Dernburg 2015)), 0.08% tricaine (Ethyl 3-aminobenzoate methanesulfonate; Sigma-Aldrich, cat. no. E10521-50G), 0.008% tetramisole hydrochloride (Sigma-Aldrich, cat. no. T1512-10G) and either 1mM or 10mM auxin (Naphthaleneacetic Acid (K-NAA), PhytoTechnology Laboratories, cat no. N610) (Martinez *et al*. 2020)) on a 22x40mm (no. 1.5) coverslip. (note: we found that poly-lysine treating the coverslips was not necessary for immobilization of the worms in most cases as long as the liquid under the agarose pad is minimal.) 7-9% agarose pads (Invitrogen, cat no. 16500500) were gently placed over the top of the worms and excess liquid was wicked away using Whatman paper. A microscope slide was adhered to the agarose pad worm coverslip sandwich using a ring of Vaseline around the pad. Worms were then imaged using the setting described below. For both SC intensity and photobleaching studies, worms were imaged immediately following mounting and worms were only kept mounted for a max of 1 hour even though worms can survive being mounted for 2-3hrs (Cahoon and Libuda 2021).

None of the worms for the studies in this paper were placed at 25ºC overnight as is commonly done to enhance expression of the fluorescently tagged proteins (Rog and Dernburg 2015; Pattabiraman *et al*. 2017). We found that in recombination deficient mutants, such as *cosa-1*, the elevated levels of GFP::SYP-2 protein in the SC with the elevation in protein expression that comes at 25ºC caused large aggregates to form in mid/late pachytene that would persist into diakinesis (Figure S8). When the worms were only grown at 20ºC without any 25ºC upshift, these aggregates did not form. Therefore, for all the experiments in this paper none of the worms were upshifted to 25ºC overnight prior to live imaging or immunofluorescence labeling.

All live imaging studies of SYP-2::GFP and mCherry::SYP-3 were imaged on a Nikon CSU SoRa Spinning Disk Microscope with a 60x water lens/N.A. 1.2 using a Z-step size of 0.3µm. For SC intensity quantifications, the laser power and exposure times were kept consistent for all genotypes. All GFP::SYP-2 imaging used the 488 laser at 16% power and 500msec exposure time. All mCherry::SYP-3 imaging used the 561 laser at 25% power and 700msec exposure time. Additionally, only the bottom half of the germline closest to the coverslip was imaged and germlines were not imaged if the position of the mounted worm caused the gut to cover germline or moved parts of the germline deeper into the worm.

The fluorescent recovery after photobleaching (FRAP) studies were performed as described in (Pattabiraman *et al*. 2017) with minor changes. Briefly, a Z-stack was taken prior to photobleaching to obtain a pre-bleach image. Then, a region of interest defined by the point tool in Elements was photobleached. A timelapse was started immediately post-photobleaching with images captured every 5 minutes for 35 minutes to monitor the fluorescence recovery. For photobleaching small regions GFP::SYP-2 or mCherry::SYP-3, a 405 laser was used with 1-5% laser power and 10-30msec exposure depending on the germline location of the nucleus and the tagged protein with GFP::SYP-2 requiring less laser power to photobleach than mCherry::SYP-3. Similar to (Pattabiraman *et al*. 2017), we also excluded adding 25mM serotonin to the live imaging media allowing for better tracking of the photobleached SC region within each nucleus during the recovery timelapse.

Immunofluorescence slides of fixed gonad were imaged on a GE DeltaVision microscope with a 63x/N.A. 1.42 lens and 1.5x optivar at 1024x1024 pixel dimensions. Images were acquired using 0.2µm Z-step size and deconvolved with softWoRx deconvolution software.

### Immunohistochemisty

Immunofluorescence was performed as described in (Libuda *et al*. 2013; Cahoon and Libuda 2021). Briefly, gonads were dissected in egg buffer with 0.1% Tween20 on to VWR Superfrost Plus slides from 18-24 hour post L4 worms. Dissected gonads were fixed in 5% paraformaldehyde for 5 minutes, flash frozen in liquid nitrogen, and fixed for 1 minute in 100% methanol at -20ºC. Slides were washed three times in PBS+0.1% Tween20 (PBST) for 5 minutes each and incubated in block (0.7% bovine serum albumin in PBST) for 1 hour. Primary antibodies (chicken anti-RAD-51, 1:1500 (Kurhanewicz *et al*. 2020; Toraason *et al*. 2021); rabbit anti-SYP-1,1:1000 (gift from Silva Lab); rabbit anti-DSB-2, 1:5000; rabbit anti-OLLAS 1:1000 (Genscript, A01658)) were added and incubated overnight in a humid chamber with a parafilm cover. Slides were then washed three times in PBST for 10 minutes each and incubated with secondary antibodies (goat anti-rabbit AlexaFluor488, ThermoFisher, cat. no. A11034; goat anti-chicken AlexaFluor488, ThermoFisher, cat. no. A11039; goat anti-rabbit AlexaFluor555, ThermoFisher, cat. no. A21428; GFP booster, Chromotek, gb2AF488-50) at 1:200 dilution for 2 hours in a humid chamber with a parafilm cover. Slides were washed two times in PBST then incubated with 2µg/mL DAPI for 15-20 minutes in a humid chamber. Prior to mounting slides were washed once more in PBST for 10 minutes and mounted using Vectashield with a 22x22mm coverslip (no. 1.5). Slides were sealed with nail polish and stored at 4ºC until imaged. GFP::MSH-5 slides were imaged within 24-48hrs of mounting due to significant signal loss in the GFP::MSH-5 staining if the slide were stored longer.

### Image Analysis and Quantification

#### FRAP quantification

The quantification of fluorescence recovery of GFP::SYP-2 and mCherry::SYP-3 was determined using FIJI. All photobleaching movies were first stabilized using the FIJI plugin “StackRegJ” (https://research.stowers.org/imagejplugins/) to reduce the nuclear and worm motion in the germline. Photobleached nuclei were cropped to exclude as much extra z volume outside the size of the nucleus as possible. Then, these nuclei were sum intensity z-projected and the fluorescence intensity of the photobleached region was monitored using the rectangle tool. A small box was drawn on the segment of SC that will be photobleached and through each frame of the timelapse the fluorescence intensity was recorded to obtain pre-bleach, bleach, and post-bleach fluorescence intensity values for a total of 35 minutes. Similar to (Pattabiraman *et al*. 2017), we also excluded any nucleus that rotated or shifted in such a way that the photobleached SC segment could not be tracked between frames of the timelapse.

Nuclei in early pachytene were defined by being within the first 5-6 rows of pachytene and nuclei in late pachytene were defined by being within the last 5-6 rows of pachytene. Mid pachytene nuclei were selected by being located within the middle region of pachytene. At each of these regions, the fluorescent intensity of three background regions of interest were determined per germline and averaged together to give the average background intensity. The average background intensity was subtracted from the fluorescence intensity of the photobleached SC segment. Additionally, the FRAP data from each SC segment was normalized such that the segment intensity pre-photobleach was 1 and the intensity immediately post-photobleached was 0. This allows us to determine the fraction of fluorescence intensity of each SYP protein that recovered following 35 minutes post-photobleaching. For oocytes, 3-5 germlines were used for GFP::SYP-2 and mCherry::SYP-3 analysis. For spermatocytes 3-6, germlines were used for GFP::SYP-2 and mCherry::SYP-3 analysis. For both sexes, 8-11 nuclei were analyzed in each region of pachytene and the specific n values in each region are reported in the figure legend. All images have been sum intensity projected and slightly adjusted for brightness and contrast. Additionally, any brightness and contrast adjustments made to oocyte images was also applied to spermatocyte images.

#### SC intensity quantification

The quantification of GFP::SYP-2 and mCherry::SYP-3 was performed using Imaris (Oxford Instruments) in combination with our whole gonad analysis, described in (Toraason *et al*. 2021). Either GFP::SYP-2 or mCherry::SYP-3 signal was surfaced in Imaris to get the sum intensity and volume of each nucleus. The start of pachytene was defined by the first row that did not contain more the 1-2 nuclei of transition zone nuclei (nuclei with DNA in a polarized or “crescent” shape morphology) and full length SC. We were able to assess the shape of the nuclei by the fluorescent hazy produced from either GFP::SYP-2 or mCherry::SYP-3 as the SC assembles. The end of pachytene was defined by the last row that contained all pachytene nuclei with the occasional single diplotene nucleus. The pachytene region was then equally divided into three zones based the length of this region within the germline to generate early pachytene, mid pachytene, and late pachytene. These criteria were used for establishing the early pachytene, mid pachytene, and late pachytene in both hermaphrodite and male germlines. Nuclei were excluded from the analysis if they were not in a single layer on the bottom half of the imaged germline rachis due to an intensity decrease caused by higher amounts of light scatter from being deeper in germline.

Our whole gonad analysis was used to align the GFP::SYP-2 or mCherry::SYP-3 surfaced nuclei along the germline length (Toraason *et al*. 2021). Each nucleus was then normalized by their volume to determine the normalized sum intensity of each nucleus during pachytene. The length of pachytene was also normalized per germline from 0 (early pachytene) to 1 (late pachytene). To calculate the average and standard deviation of the normalized SYP intensity of each nucleus during pachytene, we binned the data using a sliding window of 0.01. 7-12 germlines were analyzed for all genotypes and both sexes. During the course of this study, we discovered that from all images from May 2022 up to September 2022 needed to be corrected for a 15% drop in the power of the 561nm laser. This correction was applied to the sum intensity of mCherry::SYP-3 for all genotypes imaged during this time period, which included *spo-11* oocytes and spermatocytes, *syp-2*/+ oocytes and spermatocytes, *cosa-1* spermatocytes, *syp-3*/+ oocytes and spermatocytes, and wild type oocytes and spermatocytes. The number of nuclei analyzed within early, mid and late pachytene in each genotype is reported in the figure legends for each plot in Figure 2,3,4. All images have been sum intensity projected and slightly adjusted for brightness and contrast. Additionally, any brightness and contrast adjustments made to oocyte images was also applied to spermatocyte images.

#### SC length quantification

SC length measurements were determined on deconvolved DeltaVision images in Imaris using the filament tracer tool. Each chromosome within a nucleus was traced following the SYP-1 signal. If all six chromosomes could not be traced, then that nucleus was excluded from the analysis. Nuclei in early pachytene were defined by being within the first 5-6 rows of pachytene and nuclei in late pachytene were defined by being within the last 5-6 rows of pachytene. Mid pachytene nuclei were selected by being located within the middle region of pachytene. For spermatocyte nuclei, the smallest trace length was removed from the analysis because we inferred it to be the hemizygous *X* chromosome inappropriately assembled the SC (Jaramillo-Lambert and Engebrecht 2010). For oocytes, 10 nuclei in early pachytene were traced, 10 nuclei in mid pachytene were traced, and 12 nuclei in late pachytene were traced. For spermatocytes, 11 nuclei in early pachytene were traced, 12 nuclei in mid pachytene were traced, and 13 nuclei in late pachytene were traced.

#### Germline measurement quantifications of transition zone, DSB-2 staining, and SYP-1 assembly

Germline measurement quantification were performed using Imaris in combination with our whole gonad analysis protocol, described in (Toraason *et al*. 2021). Imaged gonad were stitched together using the FIJI (NIH) plugin Stitcher (Preibisch *et al*. 2009) and using the measurement tool in Imaris. The positions of points along the germline were recorded marking specific regions indicating the start of the germline at the pre-meiotic tip, start of the transition zone, start of DSB-2 staining, start of SYP-1 assembly, end of transition zone, end of DSB-2 zone, end of SYP-1 assembly, end of SYP-1 zone, end of pachytene and end of straggler DSB-2 nuclei. DAPI morphology was used to determine the start and end of the transition and pachytene. From these recorded point positions, we calculated the length of the transition zone, SYP-1 assembly end, DSB-2 zone end, DSB-2 straggler nuclei end and the length of pachytene. The start of the transition zone and pachytene was defined by the first row that did not contain more the 1-2 nuclei of either pre-meiotic nuclei (compact nuclei) or transition zone nuclei (nuclei with DNA in a polarized or “crescent” shape morphology), respectively. The end of pachytene was defined by the last row that contained all pachytene nuclei with the occasional single diplotene nucleus. The start of DSB-2 was determined by the position where the DSB-2 staining began in a majority of the nuclei within a row, and the end of DSB-2 staining was determined by the position were DSB-2 staining was largely absent from a majority of the nuclei. The end of the DSB-2 straggler nuclei was defined by the last nucleus in the germline with bright DSB-2 staining. The start of the SYP-1 assembly zone was defined by the germline position where small linear fragments of SYP-1 were observed, and the end of the SYP-1 assembly zone was defined by the germline position where all the nuclei in a row had full length SYP-1 with only 1-2 discontinuities. The end of the SYP-1 zone was determined by region where SYP-1 began to disassemble at the end of pachytene. The germline length was normalized per germline where 0 was the start of the germline at the pre-meiotic tip and 1 was the end of late pachytene. The number of germlines analyzed in each experiment is reported in the figure legends. All images have been max intensity projected and slightly adjusted for brightness and contrast.

#### RAD-51, MSH-5, COSA-1 quantification

Imaged gonad were stitched together using the FIJI (NIH) plugin Stitcher (Preibisch *et al*. 2009) and analyzed in Imaris as described in (Toraason *et al*. 2021) with minor changes. Each gonad from the start of pachytene through the end of pachytene was analyzed for RAD-51, MSH-5 or COSA-1 foci per nucleus, which was determined by DAPI morphology. The start of pachytene was defined by the first row that did not contain more the 1-2 nuclei of transition zone nuclei (nuclei with DNA in a polarized or “crescent” shape morphology). The end of pachytene was defined by the last row that contained all pachytene nuclei with the occasional single diplotene nucleus. The pachytene region was then equally divided into three zones based the length of this region within the germline to generate early pachytene, mid pachytene, and late pachytene. The length of pachytene was also normalized per germline from 0 (early pachytene) to 1 (late pachytene). These criteria were used for establishing the early pachytene, mid pachytene, and late pachytene in both hermaphrodite and male germlines. For RAD-51, MSH-5 and COSA-1 foci per nucleus, sliding window averages and standard error of the mean (SEM) were calculated using a 0.01 bin size. 7-12 germlines were analyzed for all genotypes and both sexes. The number of nuclei analyzed within early, mid and late pachytene in each genotype is reported in the figure legends for each plot in Figure 6. All images have been max intensity projected and slightly adjusted for brightness and contrast.

### Fertility assay

To assess fertility, L4 hermaphrodite worms were placed onto new NGM plates and were transferred every 24 hours for a total of 2 days. After 3 days from removing the hermaphrodite, each plate was scored for dead eggs. Then, the following day (4 days post removing hermaphrodite) each plate was scored for living hermaphrodite and male progeny. Additionally, any progeny with mutant Unc or Dpy phenotypes were also scored for 5 hermaphrodites in each genotype. 7-10 hermaphrodites were assayed for fertility for each genotype.

### SNP recombination mapping

SNP recombination mapping of Chromosome *II* and *X* was performed as described in (Bazan and Hillers 2011) with minor changes. *syp-2(ok307)* and *syp-3(ok785)* were generated in Bristol (N2) backgrounds and we PCR confirmed that both mutant strains carried all Bristol SNPs for both chromosomes assayed prior to mapping recombination. To generate Bristol/Hawaiian hybrids for mapping recombination, we crossed Bristol, *syp-2(ok307)* (DLW188), and *syp-3(ok785)* (DLW190) hermaphrodites to Hawaiian (CB4856) males. 8-10 Bristol/Hawaiian hybrid L4 hermaphrodites were picked off the cross plates for oocyte recombination mapping and 10-15 Bristol/Hawaiian hybrid L4 males were picked off the cross plates for spermatocyte recombination. For oocyte recombination mapping, hybrid L4 hermaphrodites of each genotype were crossed to Bristol males and male progeny were picked into 96-well plates for lysis and recombination PCR screening. For spermatocyte recombination mapping, hybrid L4 males of each genotype were crossed to Bristol hermaphrodites and male progeny were picked into 96-well plates for lysis and recombination PCR screening.

We used previously designed PCR primers and restriction digests to map six Bristol and Hawaiian SNPs on both Chromosomes *II* and *X* (Bazan and Hillers 2011). However, we were unable to get the primers to work for SNP E on Chromosome *II*, so we redesigned new primers for this SNP that worked with the existing restriction digest for SNP identification at this genomic location. All SNP positions, PCR primers, restriction digests, and band sizes of the products for the Bristol or Hawaiian SNPs can be found in Table S4. The recombination frequency or map length in centiMorgans (cM) was calculated by taking the total number of crossovers identified in each interval divided by the total chromosomes scored multiplied by 100.

### Statistics

Statistically analysis of the FRAP data, SC length, SNP recombination mapping and fertility assays was done using Prism. Mann-Whitney U tests adjusted for multiple comparisons using the Bonferroni-Dunn method was performed on the FRAP data. Kruskal-Wallis tests were performed on the SC lengths with corrections for multiple comparisons. Chi-squared tests were performed on the entire SNP recombination mapping distribution. Pairwise comparisons between recombination intervals were performed using Fisher’s Exact test. Chi-squared and Fisher’s Exact tests were performed on the fertility assays. Mann-Whitney U tests adjusted for multiple comparisons using the Bonferroni-Dunn method were performed on the SC intensity, quantification of RAD-51, MSH-5 and COSA-1, transition zone length, pachytene length, end of SYP-1 assembly zone, end of DSB-2 zone, and end of DSB-2 straggler zone using R. Each test used is indicated in the Results section next to the reported p-value and all n values are reported in the figure legends.

### Data Availability

All strains are available upon request. Strains developed as part of this study will be available at the CGC.

## Notes

### Competing Interest Statement

The authors have declared no competing interest.

