## Supplemental Figures S1-S8, Supplmental Tables S1-S4 for "Sexual dimorphic regulation of recombination by the synaptonemal complex"

### SUPPLEMENTAL FIGURES AND TABLES

#### FIGURE S1

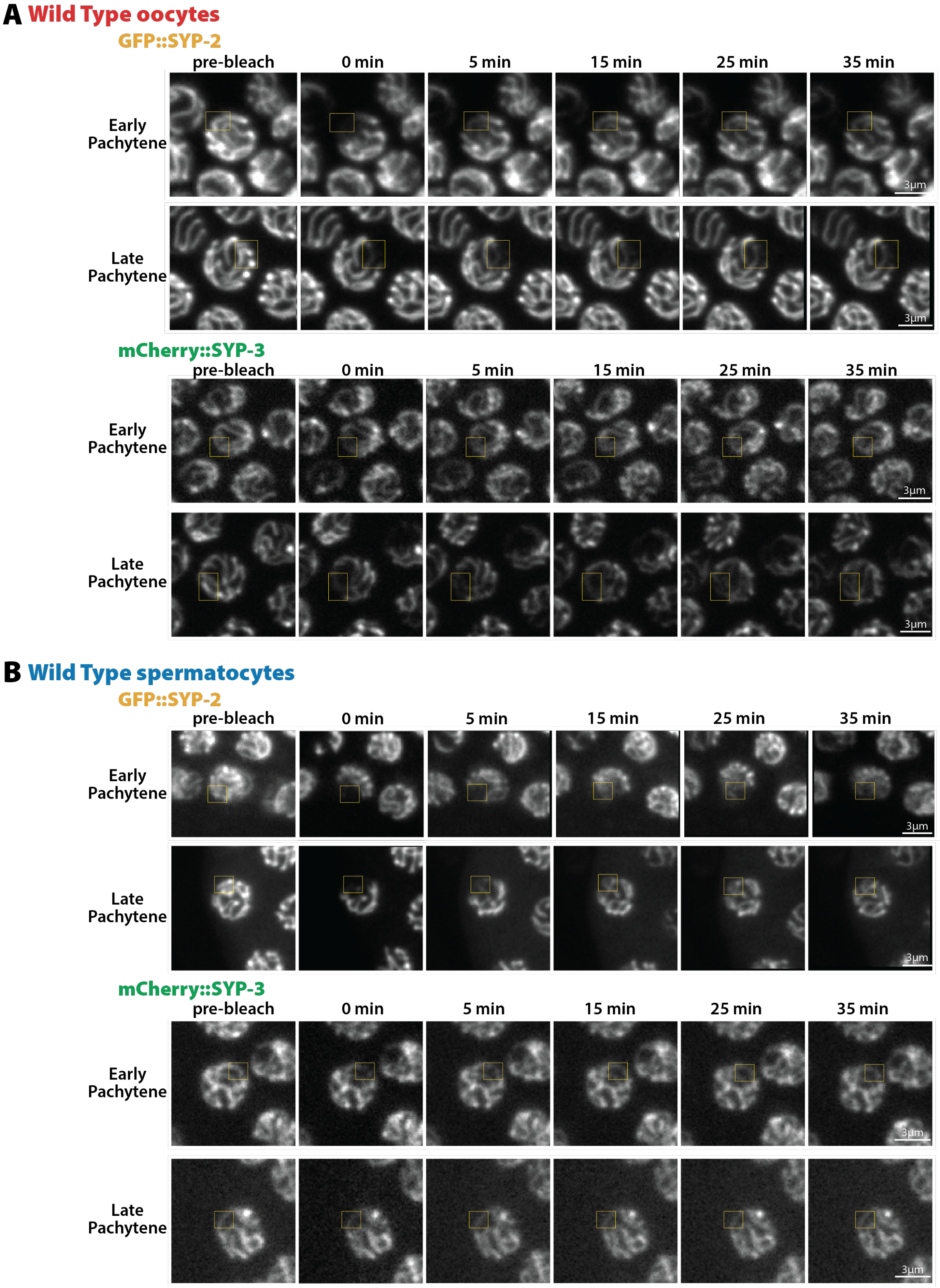

**Figure S1: Montages of FRAP from oocytes and spermatocytes.** Representative image montages of GFP::SYP-2 and mCherry::SYP-3 FRAP from oocytes **(A)** and spermatocytes **(B)** in early and late pachytene. Yellow box indicates the bleached region of each nucleus.

**FIGURE S2**

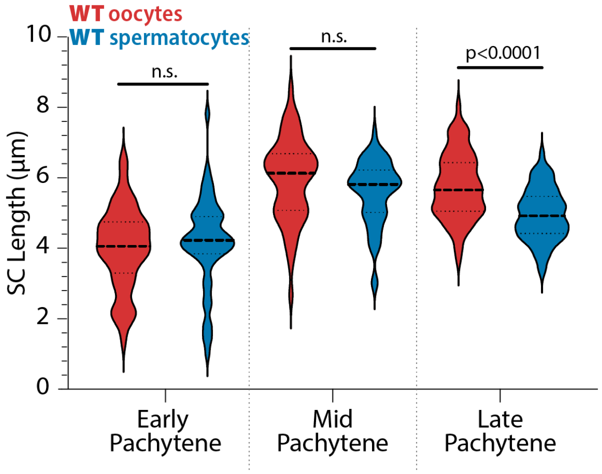


**Figure S2: SC lengths are not different between the sexes in early and mid pachytene.** Quantification of the measurement of SYP-1 length between oocytes (red) and spermatocytes (blue) throughout pachytene. Only late pachytene display significant differences in SC lengths (Mann-Whitney U test; n.s. = not significant).

### FIGURE S3

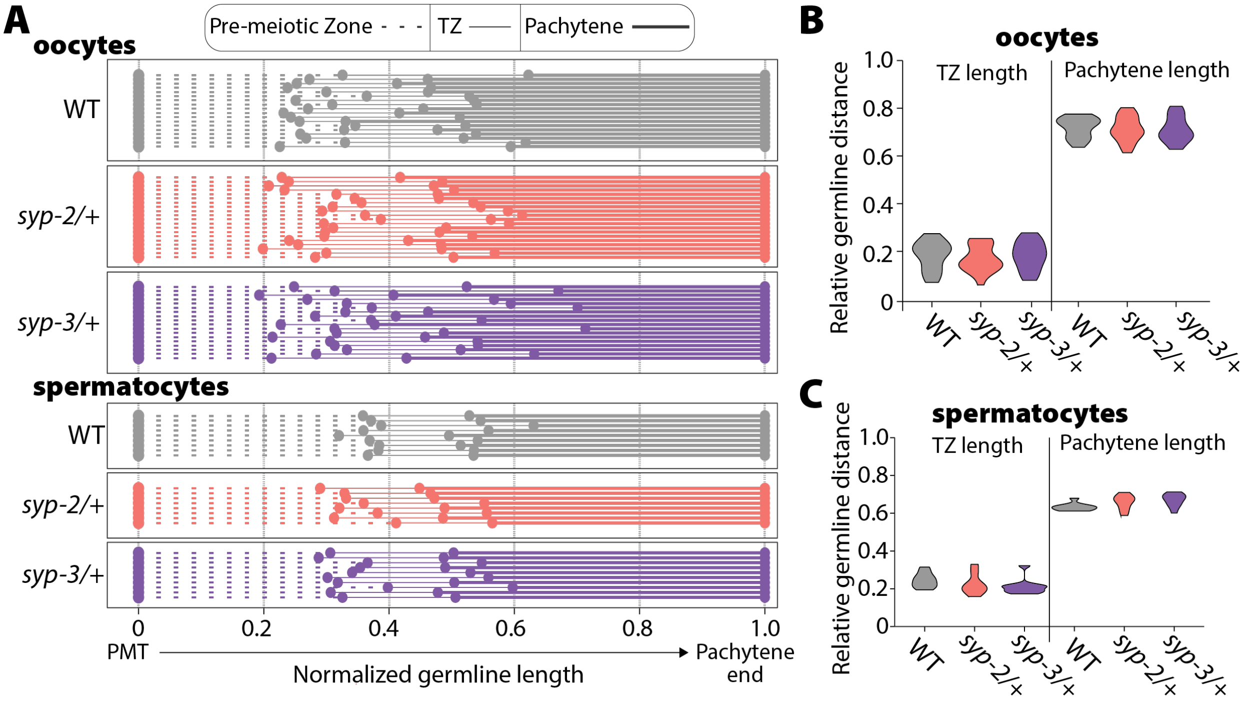


**Figure S3: Transition zone length is unaltered by SYP dosage. (A)** Measurement of the length of the pre-meiotic zone (dashed line), transition zone (thin solid line), and pachytene (thick solid line) in WT (gray), *syp-2/+* (pink), and *syp-3/+* (purple) based on DAPI morphology. All germline lengths have been normalized from the pre-meiotic tip (PMT) to the end of pachytene in both oocytes (top, saturated colors) and spermatocytes (bottom, pale colors). Each line represents an individual germline. Overall, neither *syp-2/+* nor *syp-3/+* displayed altered transition zone length. **(B,C)** Violin plots of the transition zone (TZ) length and pachytene length in WT (gray), *syp-2/+* (pink), and *syp-3/+* (purple) in oocytes **(B)** and spermatocytes **(C)**. There are no statistical differences in transition zone length or pachytene length (Mann-Whitney U test). Oocyte data is from 18 wild type germlines, 19 *syp-2/+* germlines, and 18 *syp-3/+* germlines. Spermatocyte data is from 9 wild type germlines, 8 *syp-2/+* germlines, and 10 *syp-3/+* germlines.

FIGURE S4

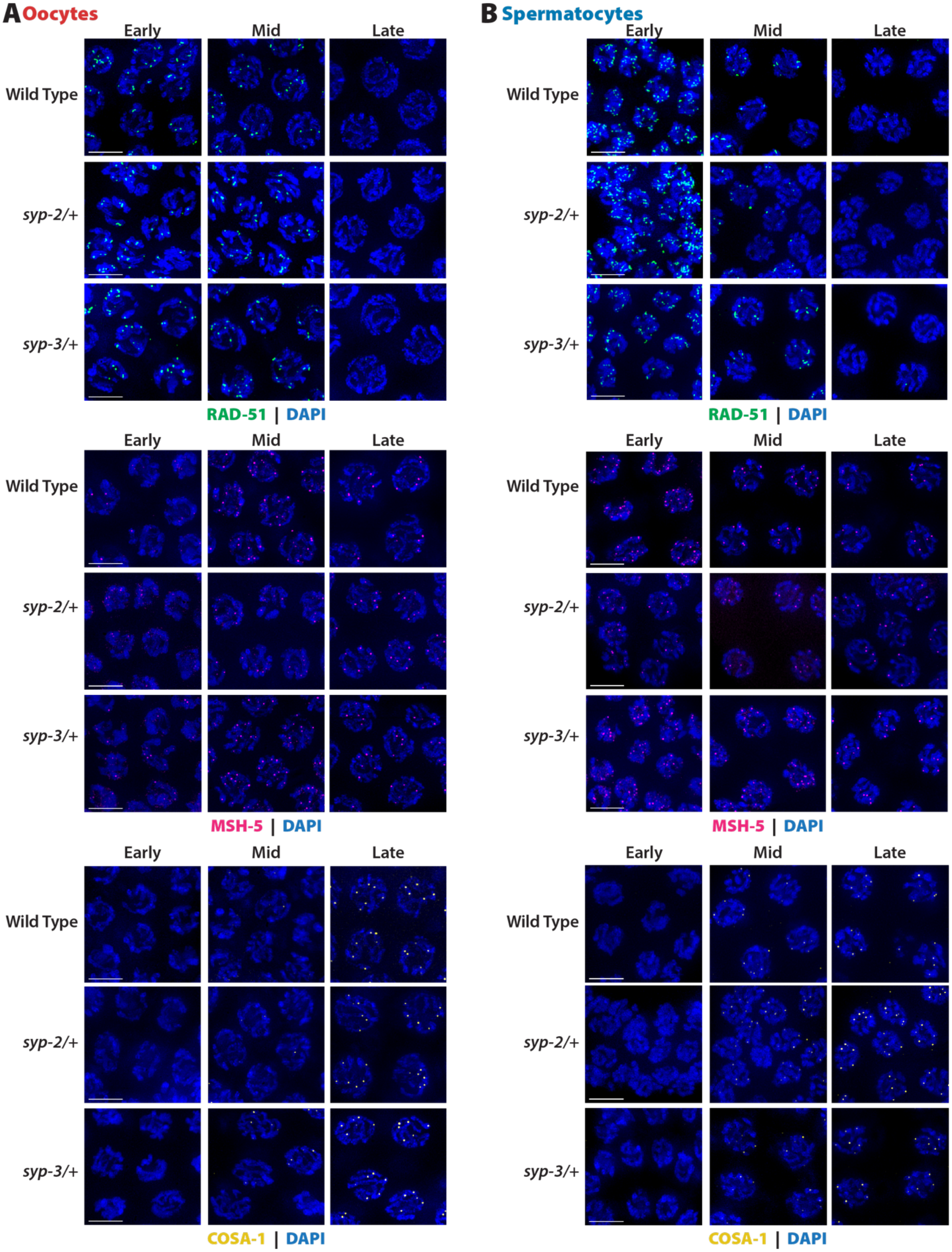


**Figure S4: Representative images of the quantification in Figure 5.** Scale bar is 5μm.

**FIGURE S5**

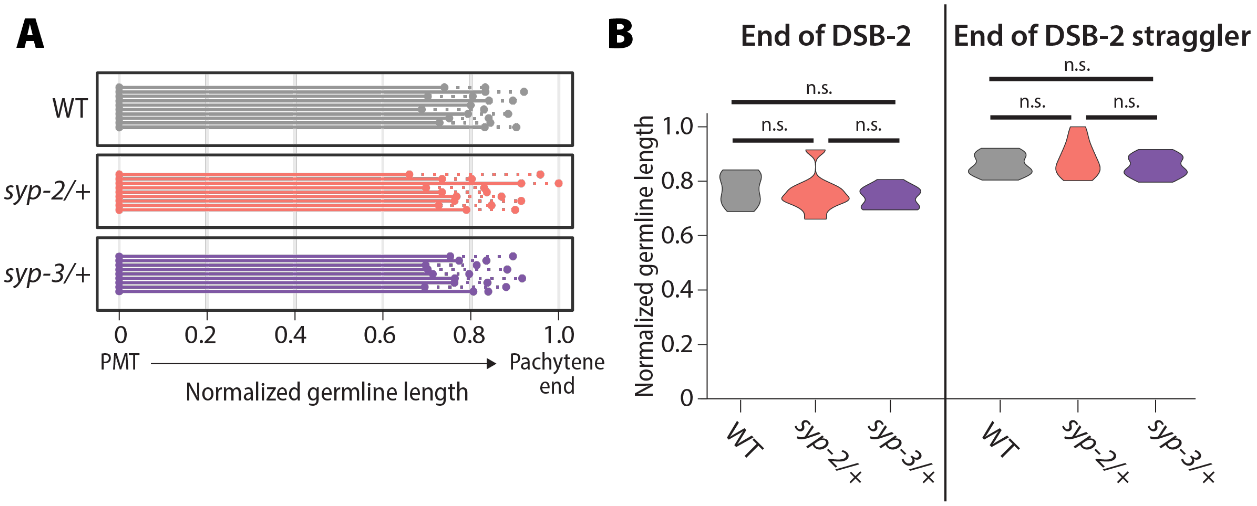


**Figure S5: DSB-2 zone is unaltered in oocytes when SYP dosage is reduced. (A)** Measurement of the length of DSB-2 staining (solid line) and the final DSB-2 stained straggler nucleus (dashed line) in WT (gray), *syp-2/+* (pink), and *syp-3/+* (purple). All germline lengths have been normalized from the pre-meiotic tip (PMT) to the end of pachytene in oocytes. Each line represents an individual germline. Overall, the zone of DSB-2 staining is largely unaltered in *syp-2/+* and *syp-3/+*. **(B)** Violin plots of the ending of the DSB-2 zone (left) and the ending of the DSB-2 straggler nuclei (right) in WT (gray), *syp-2/+* (pink), and *syp-3/+* (purple). There are no statistical differences in the end of the DSB-2 zone or the end of the DSB-2 straggler nuclei (Mann-Whitney U test). Data is from 10 wild type germlines, 9 *syp-2/+* germlines, and 9 *syp-3/+* germlines.

**FIGURE S6**

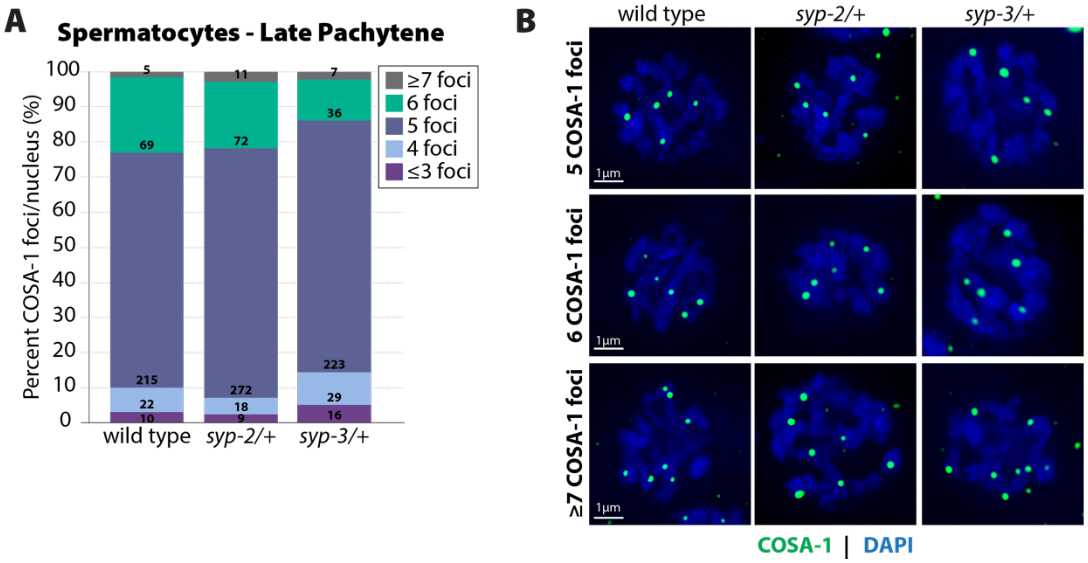


**Figure S6: Distribution of COSA-1 foci in late pachytene spermatocytes. (A)** Stacked bar plot showing the percent of nuclei with COSA-1 in each category. The individual numbers on each bar give the n value number of nuclei in each category. **(B)** Representative images of nuclei with 5, 6 and ≥7 COSA-1 foci.

FIGURE S7

**A**

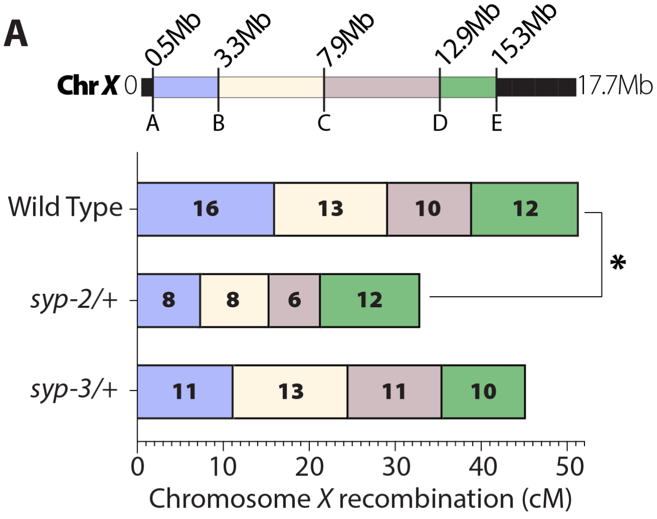


**B**

| | live herm. progeny | male progeny | dead egg | n | total mutant progeny/total non-mutant progeny | n |
|---|---|---|---|---|---|---|
| N2/CB4856 | 2484 | 1 | 483 | 10 | 4/1239 | 5 |
| *N2 syp-2/CB4856* | 1608 | 2 | 375 | 7 | 7/1111 | 5 |
| *N2 syp-3/CB4846* | 2155 | 35 | 593 | 10 | 10/1172 | 5 |
| N2/N2 | 3517 | 6 | 17 | 10 | 1/1724 | 5 |
| *syp-2/N2* | 3318 | 8 | 13 | 10 | 1/1690 | 5 |
| *syp-3/N2* | 3331 | 24 | 39 | 10 | 8/1699 | 5 |

**Figure S7: Chromosome *X* may have some chromosome distortion in *syp-2/+* oocytes.** **(A)** Recombination SNP mapping of Chromosome *X* in wild type, *syp-2/+,* and *syp-3/+* oocytes. A diagram of the 17.7Mb Chromosome *X* shows the megabase location of each SNP assayed (A-E) and the colored boxed between each SNP show the intervals where crossovers were assessed. The map length (cM) is indicated in each crossover interval. *syp-2/+* displays a significant reduction in map length on Chromosome *X* (P=0.0100, Chi-squared), while *syp-3/+* display no significant changes to wild type. The worm counts for each interval are in Table S3. **(B)** Fertility, male progeny and mutant progeny counts for wild type, *syp-2/+,* and *syp-3/+* with either all N2 (Bristol) chromosomes or N2/CB4856 (Bristol/Hawaiian). Mutant progeny were scored by displaying dumpy and/or uncoordinated mutant phenotypes. The first n from the left is for the number of worms scored with live hermaphrodite progeny, male progeny, and dead eggs. The second n from the left is for the number of worms scored having mutant progeny.

**FIGURE S8**

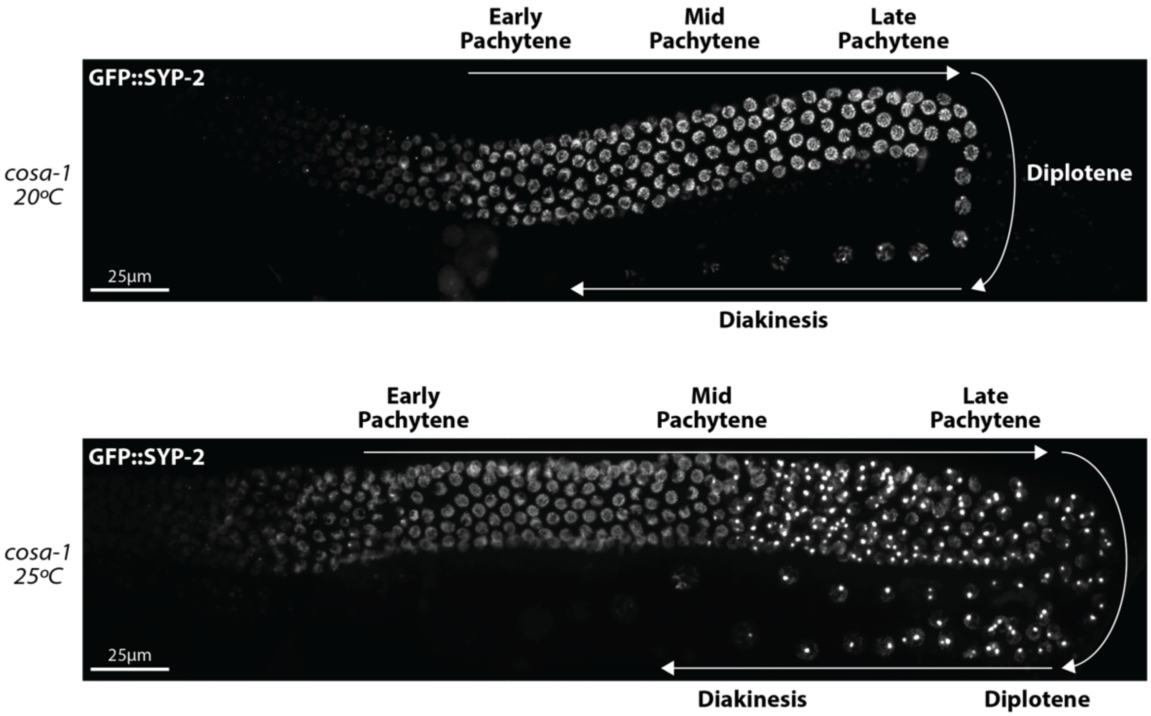


**Figure S8: GFP::SYP-2 aggregates in *cosa-1* at 25ºC during mid and late pachytene.** Representative hermaphrodite germline images of GFP::SYP-2 in *cosa-1* mutants from worms kept at 20ºC and worms upshifted to 25ºC for 18-24hrs prior to imaging. White arrows indicate the direction oocyte nuclei are moving within the germline and the stages of prophase are labeled.

**Table S1: SC intensity n values for Figures 2, 3 and 7**

| | | | Pachytene nuclei # | | | |
|---|---|---|---|---|---|---|
| Genotype | Fluorescent protein | Sex | early | mid | late | # germlines |
| WT | GFP::SYP-2 | hermaphrodite | 379 | 370 | 280 | 9 |
| WT | mCherry::SYP-3 | hermaphrodite | 294 | 490 | 411 | 14 |
| spo-11 | GFP::SYP-2 | hermaphrodite | 368 | 490 | 457 | 12 |
| spo-11 | mCherry::SYP-3 | hermaphrodite | 214 | 362 | 340 | 9 |
| cosa-1 | GFP::SYP-2 | hermaphrodite | 396 | 546 | 369 | 12 |
| cosa-1 | mCherry::SYP-3 | hermaphrodite | 535 | 814 | 553 | 17 |
| syp-2/+ | GFP::SYP-2 | hermaphrodite | 232 | 364 | 281 | 8 |
| syp-2/+ | mCherry::SYP-3 | hermaphrodite | 222 | 296 | 186 | 8 |
| syp-3/+ | GFP::SYP-2 | hermaphrodite | 301 | 323 | 255 | 10 |
| syp-3/+ | mCherry::SYP-3 | hermaphrodite | 214 | 294 | 227 | 9 |
| WT | GFP::SYP-2 | male | 227 | 240 | 220 | 12 |
| WT | mCherry::SYP-3 | male | 167 | 209 | 191 | 11 |
| spo-11 | GFP::SYP-2 | male | 178 | 204 | 155 | 9 |
| spo-11 | mCherry::SYP-3 | male | 71 | 95 | 90 | 7 |
| cosa-1 | GFP::SYP-2 | male | 131 | 186 | 157 | 7 |
| cosa-1 | mCherry::SYP-3 | male | 123 | 176 | 167 | 8 |
| syp-2/+ | GFP::SYP-2 | male | 144 | 144 | 124 | 9 |
| syp-2/+ | mCherry::SYP-3 | male | 157 | 169 | 131 | 7 |
| syp-3/+ | GFP::SYP-2 | male | 168 | 155 | 109 | 8 |
| syp-3/+ | mCherry::SYP-3 | male | 163 | 173 | 137 | 7 |

**Table S2: RAD-51, MSH-5, and COSA-1 n values for Figure 5**

| | | | Pachytene nuclei # | | | |
|---|---|---|---|---|---|---|
| **Genotype** | **Protein** | **Sex** | **early** | **mid** | **late** | **# germlines** |
| WT | RAD-51 | hermaphrodite | 375 | 412 | 339 | 8 |
| WT | MSH-5 | hermaphrodite | 465 | 618 | 519 | 11 |
| WT | COSA-1 | hermaphrodite | 533 | 459 | 384 | 9 |
| syp-2/+ | RAD-51 | hermaphrodite | 440 | 393 | 375 | 9 |
| syp-2/+ | MSH-5 | hermaphrodite | 479 | 440 | 384 | 10 |
| syp-2/+ | COSA-1 | hermaphrodite | 439 | 424 | 372 | 9 |
| syp-3/+ | RAD-51 | hermaphrodite | 382 | 336 | 282 | 9 |
| syp-3/+ | MSH-5 | hermaphrodite | 439 | 355 | 285 | 9 |
| syp-3/+ | COSA-1 | hermaphrodite | 386 | 359 | 329 | 9 |
| WT | RAD-51 | male | 471 | 420 | 476 | 10 |
| WT | MSH-5 | male | 301 | 285 | 363 | 9 |
| WT | COSA-1 | male | 303 | 269 | 301 | 12 |
| syp-2/+ | RAD-51 | male | 418 | 434 | 451 | 12 |
| syp-2/+ | MSH-5 | male | 268 | 303 | 343 | 9 |
| syp-2/+ | COSA-1 | male | 330 | 345 | 339 | 8 |
| syp-3/+ | RAD-51 | male | 376 | 296 | 383 | 10 |
| syp-3/+ | MSH-5 | male | 267 | 290 | 372 | 9 |
| syp-3/+ | COSA-1 | male | 290 | 359 | 272 | 7 |

**Table S3: Chromosome *X* SNP mapping recombination**

| | | Recombinant Intervals | | | | | |
|---|---|---|---|---|---|---|---|
| **Sex** | **Genotype** | **A—B** | **B—C** | **C—D** | **D—E** | **Non-recombinant** | **Total worms** |
| **Oocytes** | Wild Type | 43 | 35 | 28 | 34 | 129 | 269 |
| | *syp-2/+* | 22 | 22 | 18 | 29 | 172 | 263 |
| | *syp-3/+* | 28 | 35 | 29 | 27 | 134 | 253 |

**Table S4: Primers and SNP positions for recombination mapping of Chromosome *II* and *X***

| Chr | SNP | Genetic Map Position | Physical Base Pair Position | Primer Sequence (5'→3') | Restriction Enzyme | Bristol (N2) Restriction Fragments (bp) | Hawaiian (CB4856) Restriction Fragments (bp) |
|---|---|---|---|---|---|---|---|
| X | A | -19 | 0.5Mb | forward: GGTATACCGATCCCTTCAACAAG<br>reverse: TGGCAAAACACA TCCCTGTG | BspHI | 208, 156 | 364 |
| X | B | -11.1 | 3.3Mb | forward: TCGTGGCACCATAAAAGTG<br>reverse: GATTCAGATCAAACAGAGGTGG | DraI | 243 | 128, 115 |
| X | D | -0.76 | 7.9Mb | forward: GGTTCCTGGACGATAACGATGTGG<br>reverse: TCTCTCCTCTCTTTCCTCCCATTCAATC | EcoRI | 540, 228 | 768 |
| X | D | 10.1 | 12.9Mb | forward: GGCTCTGAGAAACCAACAAG<br>reverse: TGTTTGCGATGACGTGTCAG | Sau3AI | 318, 149 | 467 |
| X | E | 20.8 | 15.3Mb | forward: CGAGCAGAGATGCAGAGTTCTCAACTG<br>reverse: CGACCTGAAAGATGTGAGGTTCCTTATC | HaeIII | 280, 300 | 580 |
| II | A | -17.9 | 0.15Mb | forward: CGGAGATAGTCTCGTGGTACTG<br>reverse: CAGTCATGCTCCAAACATTCTC | DraI | 336, 93 | 288, 93, 48 |
| II | B | -4 | 4.5Mb | forward: TCATCTGTCGAGTGCTTTTG<br>reverse: CGATCGCTCAAATGGTTG | TaqI | 291, 81, 80 | 210, 81, 80, 70 |
| II | C | 3.3 | 11.1Mb | forward: TTCTCACAACTTCTTTTCCAAG<br>reverse: TTCACTATTTCCCTCGCTGG | TaqI | 572, 112, 15 | 382, 190, 112, 15 |
| II | D | 13.6 | 12.9Mb | forward: TAGGAAAGTTGTGTCCACCTGG<br>reverse: TGATGACTCCTTCTTCAGCTGC | HinfI | 449 | 288, 160 |
| II | E | 20.9 | 13.8Mb | forward: CGAGGAGATCCCACGGATTC<br>reverse: CATCGTCGTTGGCCCAGATC | TaqI | 482 | 340, 142 |
